# wQFM-TREE: highly accurate and scalable quartet-based species tree inference from gene trees

**DOI:** 10.1101/2024.07.30.605630

**Authors:** Abdur Rafi, Ahmed Mahir Sultan Rumi, Sheikh Azizul Hakim, Sohaib, Md. Toki Tahmid, Rabib Jahin Ibn Momin, Tanjeem Azwad Zaman, Rezwana Reaz, Md. Shamsuzzoha Bayzid

**Author notes:** These authors contributed equally to this work (Abdur Rafi, Ahmed Mahir Sultan Rumi).

## Abstract

Summary methods are becoming increasingly popular for species tree estimation from multi-locus data in the presence of gene tree discordance. ASTRAL, a leading method in this class, solves the Maximum Quartet Support Species Tree problem within a constrained solution space constructed from the input gene trees. In contrast, alternative heuristics such as wQFM and wQMC operate by taking a set of weighted quartets as input and employ a divide-and-conquer strategy to construct the species tree. Recent studies showed wQFM to be more accurate than ASTRAL and wQMC, though its scalability is hindered by the computational demands of explicitly generating and weighting Θ(*n*^4^) quartets. Here, we introduce wQFM-TREE, a novel summary method that enhances wQFM by circumventing the need for explicit quartet generation and weighting, thereby enabling its application to large datasets. Unlike wQFM, wQFM-TREE can also handle polytomies. Extensive simulations under diverse and challenging model conditions, with hundreds or thousands of taxa and genes, consistently demonstrate that wQFM-TREE matches or improves upon the accuracy of ASTRAL. Specifically, wQFM-TREE outperformed ASTRAL in 25 of 27 model conditions analyzed in this study involving 200-1000 taxa, with statistically significant differences in 20 of these conditions. Moreover, we applied wQFM-TREE to re-analyze the green plant dataset from the One Thousand Plant Transcriptomes Initiative. Its remarkable accuracy and scalability position wQFM-TREE as a highly competitive alternative to leading methods in the field. Additionally, the algorithmic and combinatorial innovations introduced in this study will benefit various quartet-based computations, advancing the state-of-the-art in phylogenetic estimations.

## 1 Introduction

Inferring species trees from genes sampled throughout the whole genome is a fundamental problem in molecular evolutionary biology. However, this task is complicated by the phenomenon of gene tree discordance (or heterogeneity), suggesting that different parts of the genome may have different evolutionary histories due to various biological processes, including incomplete lineage sorting (ILS), gene duplication and loss (GDL), and horizontal gene transfer (HGT) [Maddison, 1997, Warnow, 2017].

In the presence of gene tree heterogeneity, standard methods for estimating species trees, such as concatenation (which concatenates multiple sequence alignments of different genes into a single super-alignment and then estimates a tree from this alignment) can be statistically inconsistent [Degnan et al., 2009, Roch and Steel, 2015], and produce incorrect trees with high support [Kubatko and Degnan, 2007]. Therefore, “summary methods”, which operate by computing gene trees from different loci and then combining the inferred gene trees into a species tree, are becoming increasingly popular, and many of them are provably statistically consistent [Avni et al., 2015, Bayzid and Warnow, 2012, Chifman and Kubatko, 2014, Islam et al., 2020, Mahbub et al., 2021, Mim et al., 2023, Mirarab et al., 2014b, Reaz et al., 2014, Snir and Rao, 2010, Zhang, 2011].

Quartet-based summary methods have gained substantial attention as quartets (4-leaf unrooted gene trees) do not contain the “anomaly zone” [Degnan, 2013, Degnan and Rosenberg, 2006, 2009], a condition where the most probable gene tree topology may not be identical to the species tree topology. ASTRAL, the most popular summary method, tries to solve the Maximum Quartet Support Species Tree (MQSST) problem [Mirarab et al., 2014b]. It takes a set of gene trees as input and seeks to find a species tree so that the number of induced quartets in the gene trees that are consistent with the species tree is maximized. wQFM and wQMC are alternative heuristics for solving the MQSST problem, which involve deducing individual quartets and then combining them into a cohesive species tree in a divide-and-conquer fashion. Since they have to preprocess and analyze Θ(*n*^4^) quartets induced by a set of gene trees on *n* taxa, the running time becomes prohibitively large with increasing numbers of taxa. Consequently, although wQFM was shown to have consistently outperformed wQMC and ASTRAL through extensive simulation studies [Mahbub et al., 2021, 2022], even when gene trees are incomplete [Mahbub et al., 2022], it failed to gain widespread attention from the systematists. Han and Molloy have recently developed TREE-QMC, a substantially faster version of wQMC, eliminating the need for explicitly computing the weighted quartet distribution displayed by the input gene trees [Han and Molloy, 2023]. Mim et al. [2023] have recently presented a fast implementation of the original Quartet Fiduccia–Mattheyses (QFM) algorithm [Reaz et al., 2014], which solves the unweighted version of the maximum quartet consistency problem. In this work, we introduce a novel method, wQFM-TREE (wQFM applied directly to gene trees), employing an innovative technique to conduct the same heuristic search as wQFM, which obviates the need to decompose the input gene trees into their induced quartets. With a time complexity of *O*(*n*^3^*k* log *n*) under certain assumptions (n is the number of taxa and k is the number of gene trees), wQFM-TREE substantially improves upon the running time of wQFM, making it scalable to datasets with thousands of taxa and genes. Furthermore, similar to the findings reported in previous studies [Han and Molloy, 2023, Mahbub et al., 2021, 2022] for small to moderate-sized datasets, wQFM-TREE demonstrated excellent accuracies on large datasets and outperformed ASTRAL on most of the datasets analyzed in this study.

## 2 Materials and Methods

We begin with a brief overview of the original wQFM. We next describe our proposed techniques introduced in wQFM-TREE to make it scalable.

### 2.1 Background on wQFM

Given a set 𝒢 = {*g*_1_, *g*_2_, …, *g*_*k*_} of *k* gene trees on taxa set *𝒳*, wQFM computes weights for every possible quartet *ab*|*cd*, where *ab*|*cd* denotes the unrooted quartet tree with leaf set *a, b, c, d* ∈ *𝒳* in which the pair *a, b* is separated from the pair *c, d* by an edge. Next, wQFM constructs a species tree using a divide-and-conquer algorithm, which operates in the following top-down manner: a good non-trivial bipartition of *𝒳* is produced using the weighted quartets, rooted trees are calculated on each part of the bipartition by running wQFM recursively, and then these rooted trees are combined into a rooted tree on the full taxa set (Supplementary Figure 1).

Central to wQFM’s search algorithm is generating 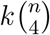 quartets induced by the input gene trees, computing the weights of these induced quartets based on gene tree frequency, and scoring a candidate bipartition based on the weights of the input quartets that are consistent and conflicting with that particular bipartition. Generating weighted quartets (and saving them to a file using appropriate I/O operations) is an extremely time and memory-consuming step and constitutes a significant portion of wQFM’s overall runtime. This weighted quartet generation step has actually been the main limitation for the broader application of wQFM to larger datasets.

Each divide step of the wQFM algorithm splits the set of taxa into two disjoint subsets (a bipartition of the taxa set, each representing a subproblem). A unique “dummy taxon” which is an artificial taxon is added to both of the subsets so that the solutions to the subproblems can be combined within a divide-and-conquer framework (Supplementary Figure 1). The technique to find a bipartition begins with an initial bipartition, followed by a heuristic iterative strategy to improve it. Within this iterative process, each iteration begins with the bipartition from the previous iteration and tries to improve it by moving taxa from one partition to another. The iterative improvement strategy is described in detail in [Reaz et al., 2014]. The effectiveness of such a move, which we call “Gain”, is quantified by the differences in bipartition scores before and after a taxon transfer. Mim et al. [2023] showed that given Θ(*n*^4^) quartets, one iteration of this technique of improving an initial bipartition takes *O*(*n*^4^) time which is very high. Producing an initial bipartition at every bipartitioning step is also a highly time-consuming process. The computation of the initial bipartition entails working with sorted weighted quartets, aiming to identify a bipartition consistent with quartets of substantial weights. Sorting Θ(*n*^4^) weighted quartets and selecting a good initial bipartition accordingly is a highly time-consuming step.

### 2.2 Overview of wQFM-TREE

wQFM-TREE has incorporated new algorithms and features to eliminate the necessity of generating and analyzing all potential quartets, employing the following two key strategies: i) using a gene tree consensus-based method to compute an initial bipartition for each divide step and ii) utilizing combinatorial and graph theoretic techniques to compute the scores of candidate bipartitions directly from gene trees, thereby eliminating the need for generating and computing the weights of the induced quartets. Both of these strategies heavily depend on a certain tree structure that we define for dummy taxa (discussed in Section 2.3) in our proposed method. One iteration of the iterative improvement strategy for finding a bipartition takes *O*(*n*^2^*k* log *n*) with our proposed techniques instead of the *O*(*n*^4^) runtime of the original wQFM, with certain assumptions on subproblem sizes (details in Supplementary Section 3).

### 2.3 Tree structure of a dummy taxon and weighting scheme

In our divide step, we create a bipartition (*A, B*) of the taxa set and create two new subproblems, one with taxa set *A*∪{*X*_1_} and the other with taxa set *B*∪{*X*_2_}, where *X*_1_ and *X*_2_ are dummy taxa. Note that, unlike the original wQFM, we differentiate between the dummy taxa added to the two subproblems. *X*_1_ essentially represents the taxa in *B*, and *X*_2_ represents the taxa in *A*. We model *X*_1_ and *X*_2_ as rooted tree structures, where the children of the roots are the taxa in *B* and *A*, respectively. If *A, B* contain any dummy taxon, they will appear as subtrees in the tree structure of *X*_2_, *X*_1_ respectively, resulting in a recursive tree structure (Figure 1). Therefore, the tree representation of a dummy taxon *X* is a rooted tree with height *>* 1. The leaves of this tree are real taxa, and any internal node is a dummy taxon which was introduced in an earlier subproblem. We say that the leaves or real taxa in the tree structure of *X* are *under dummy taxon X*, and denote this set of real taxa under *X* as *X*_*R*_.

**Figure 1:**
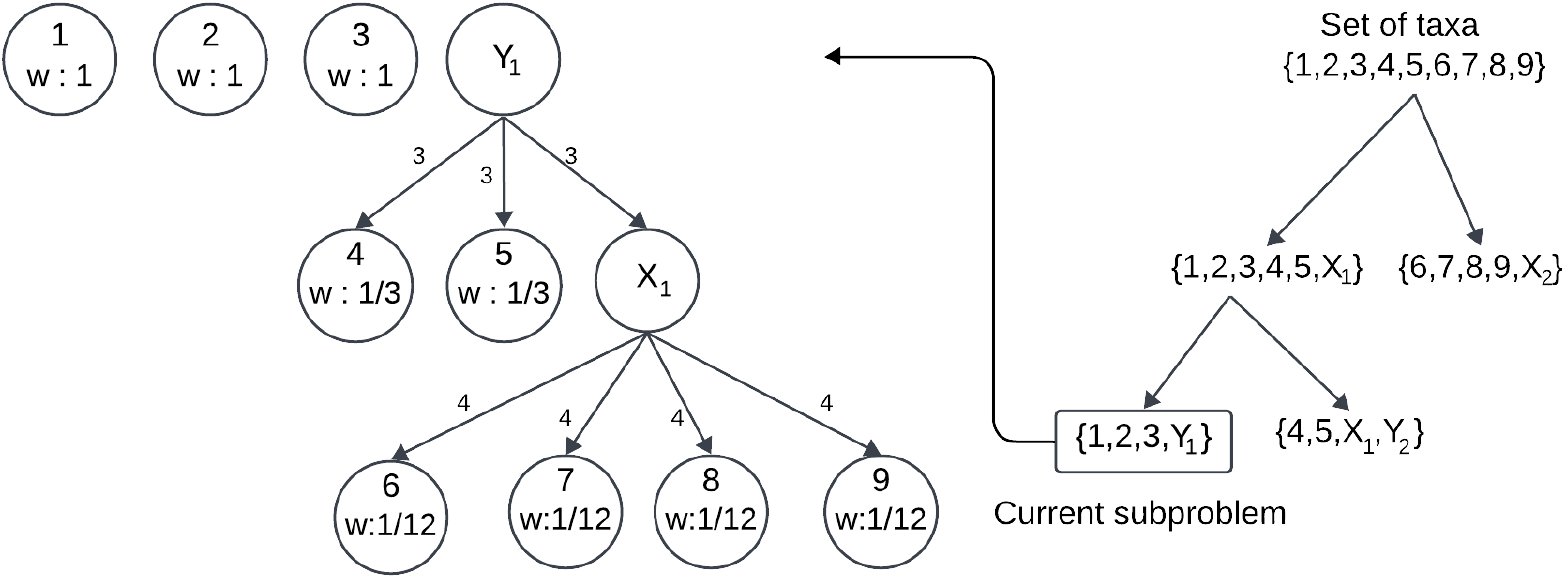
Tree structure of a dummy taxon, and the assignment of weights to real taxa under this dummy taxon. Here *Y*1 is a dummy taxon that represents {4, 5, *X*_1_}, where *X*_1_ is another dummy taxon representing {6, 7, 8, 9}. The entire tree structure (root labeled by *Y*_1_) is the representation of dummy taxon *Y*_1_. The real taxa under *Y*1 appear as leaves in the tree structure. Edge weights for each edge are shown. The weight of a leaf is the multiplicative inverse of the product of the edge weights from the root to the leaf.

Now we describe how we assign weight *w*(*a*) to a taxon *a* that is used to compute the score of a candidate bipartition as described in Section 2.5. We assign unit weight to the real taxa that are not under any dummy taxa of the current subproblem. Note that the contribution of a dummy taxon to the score of a candidate bipartition should be equal to that of a real taxon as both can be considered equivalent in a subproblem. Therefore, for a dummy taxon *X*, we assign fractional weights to the real taxa in *X*_*R*_ in such a way that 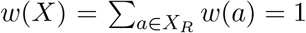. We refer to this assignment as weight normalization and we describe our motivation behind it in more detail in Supplementary Section 2.3. The normalization process is as follows. We first assign weights to the edges of the tree structure corresponding to *X*. We then use these edge weights to calculate *w*(*a*) (*a* ∈ *X*_*R*_). Refer to Figure 1 for an example. The weight of an edge (*u, v*) with *u* as the parent node, is defined as the number of children of *u*. Let *p*_*a*_ be the set of edges in the path from root to *a*. We define *w*(*a*) as 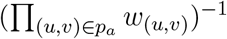. Therefore, for any real taxon *a, w*(*a*) is defined as follows.

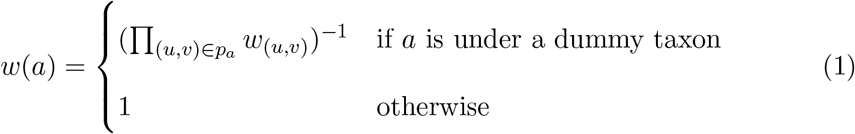

We note that a similar weighting mechanism is used in the TREE-QMC method [Han and Molloy, 2023] for weighting quartets to normalize the “Quartet Graph”. We use this weighting mechanism with a slightly different motivation than the TREE-QMC method for the following use cases: i) to construct initial bipartition from consensus tree of gene trees, and ii) to calculate the score of a candidate bipartition using weighted satisfied and violated quartets.

### 2.4 Computing an initial bipartition for a divide step

Let *S* be the set of taxa of a subproblem in a divide step of our method. We need to create an initial bipartition of *S* which will subsequently be improved in an iterative manner. To do so, we create and utilize a greedy consensus tree of the input gene trees, also known as the majority rule extended (MRE) tree, using the Paup package [Swofford, 2002]. This works even for incomplete gene trees, which are very common in real datasets.

Each bipartition (*A, B*) corresponding to each edge of the consensus tree defines a bipartition (*S*_*A*_, *S*_*B*_) of *S* as follows. *S* may contain both real and dummy taxa. The real taxa in *S* are partitioned according to (*A, B*) meaning that if a taxon *s* ∈ *A*, then *s* ∈ *S*_*A*_. We now describe how a dummy taxon *X* in *S* is assigned to a partition (*S*_*A*_ or *S*_*B*_). We compute the weight of each taxon in *X*_*R*_ (the set of real taxa present in the tree structure of *X*) as described in Section 2.3. *X*_*R*_ can be partitioned to *S*_*A*_ and *S*_*B*_ according to (*A, B*) similar to how we partition the real taxa in *S* with respect to (*A, B*). Next, we assign *X* to the partition *S*_*A*_ if the sum of weights of the real taxa in *X*_*R*_ that belong to *A* is greater than the sum of weights of the real taxa in *X*_*R*_ that belong to *B*. Otherwise, we assign *X* to the partition *S*_*B*_. Thus, we find a bipartition (*S*_*A*_, *S*_*B*_) of *S* with respect to (*A, B*) in the consensus tree. Next, we score (*S*_*A*_, *S*_*B*_) using the scoring scheme described in Section 2.5. We find and score all such bipartitions of *S* corresponding to the bipartitons in the consensus tree. We choose the bipartition of *S* with the highest score as the initial bipartition.

### 2.5 Scoring a candidate bipartition directly from gene trees

Let 𝒢 be a set of unrooted gene trees on taxa set *𝒳*. We allow the gene trees to be non-binary and incomplete (i.e., some taxa could be missing in a gene tree). Let *𝒳*^(*g*)^ be the set of real taxa in a gene tree *g* ∈ 𝒢. Let (*A, B*) be the given bipartition. Let *R*_*A*_ and *D*_*A*_ be the sets of real and dummy taxa in *A*, respectively. Let *F*_*A*_ be the set of all real taxa that are either in *R*_*A*_ or under a dummy taxon *X* ∈ *D*_*A*_, i.e., 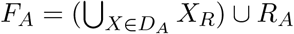. We define *R*_*B*_, *D*_*B*_, *F*_*B*_ similarly.

Our algorithm assigns weight to each real taxon in *𝒳* according to Equation 1. These weights are used to calculate the weights of quartets. Let *a, b, c, d* ∈ *𝒳*. We define the weight of an unordered pair of taxa, *p* = {*a, b*} as *w*(*p*) = *w*(*a*) · *w*(*b*). We consider a quartet *q* = *ab*|*cd* to be composed of 2 unordered pairs {*a, b*} and {*c, d*}. We define its weight *w*(*ab*|*cd*) to be *w*({*a, b*}) · *w*({*c, d*}). We define the weight of a set *Q* of quartets as *w*(*Q*) = ∑_*q*∈*Q*_ *w*(*q*). We define the weight of a set of unordered pairs similarly.

Given a bipartition (*A, B*), each quartet *ab*|*cd* where no two taxa in {*a, b, c, d*} are under the same dummy taxon of the current subproblem, can be classified as one of the following three types.

1. **Satisfied:** where either {*a, b*} ⊆ *F*_*A*_ and {*c, d*} ⊆ *F*_*B*_ or {*a, b*} ⊆ *F*_*B*_ and {*c, d*} ⊆ *F*_*A*_
2. **Deferred:** where *P* ⊆ *F*_*A*_ or *P* ⊆ *F*_*B*_, *P* ⊆ {*a, b, c, d*}, and |*P* | ≥ 3.
3. **Violated:** all other quartets.

Note that we do not consider the quartets that have at least two taxa under the same dummy taxon. This is because although a dummy taxon may have multiple real taxa under it, it is considered a single taxon for the current subproblem. Therefore, a dummy taxon should not contribute more than once in a quartet. In the original wQFM, we do not need to consider this extra condition as it enumerates all the quartets and updates the set of considered quartets for each subproblem in the divide step.

We note that if a gene tree *g* is non-binary, then for some *a, b, c, d* ∈ *𝒳*^(*g*)^, a quartet structure may not be formed/resolved (i.e., there is no branch separating two taxa from the other two). We call such quartets *unresolved*.

Let *S*^(*g*)^, *V* ^(*g*)^ be the sets of satisfied and violated resolved quartets respectively in a gene tree *g* ∈ 𝒢. Then the score *Score*(*A, B*, 𝒢) of a candidate bipartition (*A, B*) with respect to 𝒢 is defined as *Score*(*A, B*, 𝒢) = ∑_*g*∈*𝒢*_ (*w*(*S*^(*g*)^) − *w*(*V* ^(*g*)^)). Let *U* ^(*g*)^ be the set of satisfied or violated unresolved quartets. Since *S*^(*g*)^, *V* ^(*g*)^, *U* ^(*g*)^ are disjoint sets, *w*(*S*^(*g*)^ ∪ *V* ^(*g*)^ ∪ *U* ^(*g*)^) = *w*(*S*^(*g*)^) + *w*(*V* ^(*g*)^) + *w*(*U* ^(*g*)^). We can rewrite (*w*(*S*^(*g*)^) − *w*(*V* ^(*g*)^)) as (2*w*(*S*^(*g*)^) − (*w*(*S*^(*g*)^) + *w*(*V* ^(*g*)^) + *w*(*U* ^(*g*)^)) + *w*(*U* ^(*g*)^)).

Thus, we get the following equation.

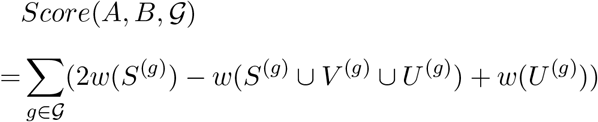

We do such restructuring of this equation for computational efficiency. Calculating *w*(*S*^(*g*)^) and *w*(*V* ^(*g*)^) separately requires traversing each internal node of the tree *g* and perform certain calculations (details in Section 2.5.1 for *w*(*S*^(*g*)^)). However, *w*(*S*^(*g*)^ ∪ *V* ^(*g*)^ ∪ *U* ^(*g*)^) can be calculated directly from *X* ^(*g*)^ (details in Section 2.5.2). Hence, by eliminating *w*(*V* ^(*g*)^) and incorporating *w*(*S*^(*g*)^ ∪ *V* ^(*g*)^ ∪ *U* ^(*g*)^) into the equation, the computation becomes notably faster in practice. Although we have to compute *w*(*U* ^(*g*)^) in this approach, it is still more efficient because calculating *w*(*U* ^(*g*)^) requires us to consider only the polytomy nodes (details in Section 2.5.3).

#### 2.5.1 Computing *w*(*S*^(*g*)^), the sum of weights of satisfied resolved quartets in a gene tree *g*

Let *u* be an internal node in *g* and *deg*(*u*) denote the degree of *u*. Removing *u* will create *deg*(*u*) components of *g*. We define *C*^(*g,u*)^ to be the set of these components and use 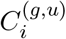, 1 ≤ *i* ≤ *deg*(*u*) to refer to the *i*^*th*^ component. We say, a real taxon 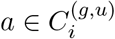 if *a* is in component 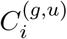.

The subtree of a fully resolved tree restricted to a quartet exhibits two degree-three nodes. We refer to these nodes as *anchors* of the quartet on that tree. Now, let *S*^(*g,u*)^ be the set of all resolved satisfied quartets *ab*|*cd* such that *a, b* ∈ *F*_*A*_, *c, d* ∈ *F*_*B*_ and *a, b* belong to two distinct components in *C*^(*g,u*)^ and both *c, d* pertain to a third component of *C*^(*g,u*)^. We observe that *S*^(*g,u*)^ actually contains the quartets, which have *u* as the anchor directly connected to *a, b* in the quartet tree. All such sets *S*^(*g,u*)^, across every internal node *u*, form a partition of *S*^(*g*)^ (Lemma 1 in Supplementary Section 2). As a result, *w*(*S*^(*g*)^) = ∑*u w*(*S*^(*g,u*)^).

To compute *w*(*S*^(*g,u*)^), we further split *S*^(*g,u*)^ into disjoint sets 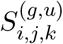, 1 ≤ *i, j, k* ≤ *deg*(*u*), *i < j, k* ≠ *i, j*. 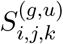 contains the satisfied quartets *ab*|*cd* where *a, b* ∈ *F*_*A*_, 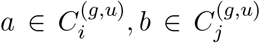, and *c, d* ∈ *F*_*B*_, *c*,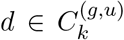. Let 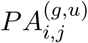 be the set of unordered pairs {*a, b*} where *a, b* are not under the same dummy taxon, *a, b* ∈ *F*_*A*_, and 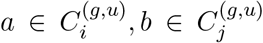. Similarly, let 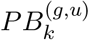 be the set of unordered pairs {*c, d*} where *c, d* are not under the same dummy taxon, *c, d* ∈ *F*_*B*_, and *c*, 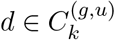, *c* ≠ *d*. Therefore, for a quartet 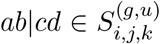, the unordered pair {*a, b*} 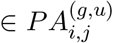 and the unordered pair {*c, d*} 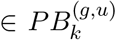 where *k* ≠ *i, j*. As a result,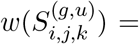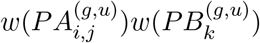. Then we compute

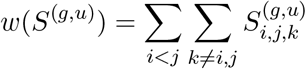

We restructure this expression for efficient implementation, which is described in Supplementary Section 2.1.3. We can calculate 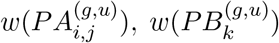 without enumerating the sets 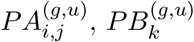 (see Supplementary Section 2.1 for details). Thus, for a given candidate bipartition (*A, B*), we can compute the weight of the satisfied resolved quartets in *g* without explicitly enumerating its induced set of quartets.

#### 2.5.2 Computing *w*(*S*^(*g*)^ ∪ *V* ^(*g*)^ ∪ *U* ^(*g*)^), the sum of weights of satisfied or violated quartets in *g*

We notice that, each quartet *q* = *ab*|*cd* ∈ (*S*^(*g*)^ ∪ *V* ^(*g*)^ ∪ *U* ^(*g*)^) has two taxa from *F*_*A*_, the other two from *F*_*B*_ and no pair of taxon from {a,b,c,d} is under the same dummy taxon. Let *PA*^(*g*)^ be the set of unordered pairs {*a, b*} where *a, b* are not under the same dummy taxon and *a, b* ∈ *F*_*A*_ ∩ *𝒳* ^(*g*)^ (the intersection with *𝒳*^(*g*)^ allows our computation to be applicable for incomplete gene trees where some taxa are missing). Similarly, let *PB*^(*g*)^ be the set of pairs {*c, d*} where *c, d* are not under the same dummy taxon and *c, d* ∈ *F*_*B*_ ∩ *𝒳* ^(*g*)^. Each pair {*a, b*} ∈ *PA*^(*g*)^ forms one quartet with each pair {*c, d*} ∈ *PB*^(*g*)^. Thus, we obtain

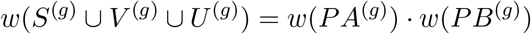

We provide the expressions for calculating *w*(*PA*^(*g*)^) and *w*(*PB*^(*g*)^) in Supplementary Section 2.1.4.

#### 2.5.3 Computing *w*(*U* ^(*g*)^), the sum of weights of unresolved satisfied or violated quartets in *g*

For each *ab*|*cd* ∈ *U* ^(*g*)^, we can find only one internal node *u* in *g* such that *a, b, c, d*, all come from 4 different components in *C*^(*g,u*)^ and *a, b* ∈ *F*_*A*_, *c, d* ∈ *F*_*B*_. Let *U* ^(*g,u*)^ be the set of these quartets. We obtain *w*(*U* ^(*g*)^) by adding *w*(*U* ^(*g,u*)^) for each polytomy node *u*.

Similar to computing *w*(*S*^(*g,u*)^), we further split *U* ^(*g,u*)^ into disjoint sets 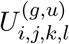, 1 ≤ *i, j, k, l* ≤ *deg*(*u*), *i < j, k < l, k* ≠ *i, j*; *l* ≠ *i, j*. 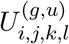 contains the unresolved quartets *ab*|*cd* of *U* ^(*g,u*)^ where 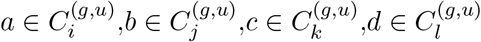. Let 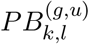 be the set of unordered pairs {*c, d*} where *c, d* are not under the same dummy taxon, *c, d* ∈ *F*_*B*_, and 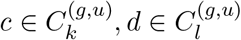. Note that each pair 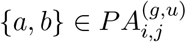 forms an unresolved quartet with each pair 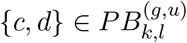 when *k < l, k* ≠ *i, j*; *l* ≠ *i, j* and these quartets form the set 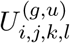. Therefore 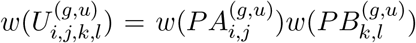.

Then

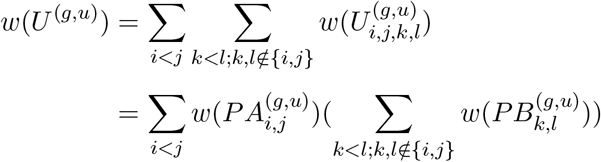

We employ this equation to derive an expression for *w*(*U* ^(*g,u*)^) that enables efficient implementation (Supplementary Section 2.1.5). We present an example showing detailed calculations of our scoring method in Supplementary Section 2.2.

#### 2.5.4 Gain calculation

Note that the wQFM algorithm improves a bipartition iteratively by transferring taxa from one partition to the opposite, and the effectiveness of a move is quantified by *Gain* which is the change of scores after transferring a taxon. The procedure for calculating gains using our scoring method is outlined in Supplementary Section 2.4.

### 2.6 Time Complexity

Assuming that the input gene trees are fully resolved or polytomies are resolved arbitrarily, and the subproblems are produced in a perfectly balanced fashion, wQFM-TREE can run in *O*(*n*^3^*k* log *n*) time, where *n* and *k* denote the number of taxa and gene trees, respectively. A detailed time complexity analysis is presented in Section 3 of the Supplementary Materials. We note that TREE-QMC has a time complexity of *O*(*n*^3^*k*) considering balanced subproblems (TREE-QMC needs to resolve polytomies arbitrarily).

## 3 Experimental studies

### 3.1 Datasets

#### Simulated dataset

We evaluated wQFM-TREE on large-scale simulated datasets from the ASTRAL-II study [Mirarab and Warnow, 2015]. We used 8 model conditions from the ASTRAL-II dataset, simulated under the Yule model from model species trees, which are characterized by three parameters: the height of the species tree, the speciation rate, and the number of taxa. In six model conditions, the number of taxa was set to 200 with varying tree lengths (500 K, 2 M, and 10 M generations) and speciation rates (1e-6 and 1e-7 per generation). The remaining two model conditions contain 500 and 1000 taxa with tree shapes fixed to 2 M/1e-06. For all the model conditions, we assessed the performance on varying the number of genes (50, 200, and 1000). Each model condition has 50 replicates of data. We analyzed all 50 replicates except for the 1000 taxa dataset, for which we analyzed 20 replicates (*R*1 − *R*21, excluding *R*8) in our study. We excluded *R*8 as wQFM-TREE failed to analyze it in the model condition with 1000 gene trees within the allotted 48 hours of computation due to presence of high amount of polytomy.

Moreover, we compared wQFM-TREE to the wQFM on the 48-taxon avian and 37-taxon mammalian simulated datasets from [Mirarab et al., 2014a] as the original wQFM cannot handle large datasets with hundreds of taxa. These datasets were simulated based on previously published species trees for 48 birds [Jarvis et al., 2014] and 37 mammals [Song et al., 2012], respectively. We also analyzed a 2000-taxon dataset from the ASTRAL-III study [Zhang et al., 2018] to test the scalability of wQFM-TREE.

#### Empirical dataset

We re-analyzed the green plant dataset [1kp, 2019] – one of the largest phylogenomic datasets analyzed so far – from the One Thousand Plant Transcriptomes Initiative. This dataset contains 410 single-copy gene family trees across 1178 species, including green plants (Viridiplantae), glaucophytes (Glaucophyta), red algae (Rhodophyta), and outgroup species.

### 3.2 Methods and measurements

We compared wQFM-TREE to the best existing species tree estimation methods wQFM, ASTRAL-III (v.5.7.8), and TREE-QMC.

On the simulated datasets, we compared the estimated trees with the model species tree using normalized Robinson-Foulds (RF) distance [Robinson and Foulds, 1981]. For the biological dataset, we compared the estimated species trees to established evolutionary relationships. We assessed support values in the estimated trees using local posterior probabilities [Sayyari and Mirarab, 2016] computed by ASTRAL. We analyzed multiple replicates of data for various model conditions and performed a two-sided Wilcoxon signed-rank test [Wilcoxon, 1945]. We set the threshold *α* = 0.05 to measure the statistical significance of the differences between two methods.

## 4 Results and discussion

### 4.1 Results on simulated datasets

We first compare wQFM-TREE with wQFM in terms of accuracy and running time. Next, we extensively compare wQFM-TREE with ASTRAL and TREE-QMC on large datasets with hundreds of taxa and genes.

#### 4.1.1 Comparing wQFM-TREE with wQFM

Due to space constraints, we provide the experimental results in Supplementary Section 4.1. These results suggest that wQFM-TREE is dramatically faster than the original wQFM without sacrificing the accuracy.

#### 4.1.2 Comparing wQFM-TREE with other coalescent-based methods

##### Results on 200-taxon dataset

The evaluation of wQFM-TREE, ASTRAL, and TREE-QMC is presented in Figure 2 for the 200-taxon datasets under various model conditions with varying tree lengths, speciation rates, and numbers of taxa. On relatively short tree lengths (i.e., 500K generations), wQFM-TREE performed comparably with ASTRAL and the differences between the methods were often statistically insignificant. wQFM-TREE performed better than ASTRAL with statistical significance in 1 model condition and so did ASTRAL in 2 model conditions. However, as the number of generations increased, wQFM-TREE demonstrated superior performance. Specifically, wQFM-TREE outperformed ASTRAL in all 12 model conditions with 2M and 10M generations, and the improvements were statistically significant (*P <* 0.05) in 11 of these 12 model conditions. Additionally, the improvements of wQFM-TREE over ASTRAL increased with increasing numbers of genes.

**Figure 2:**
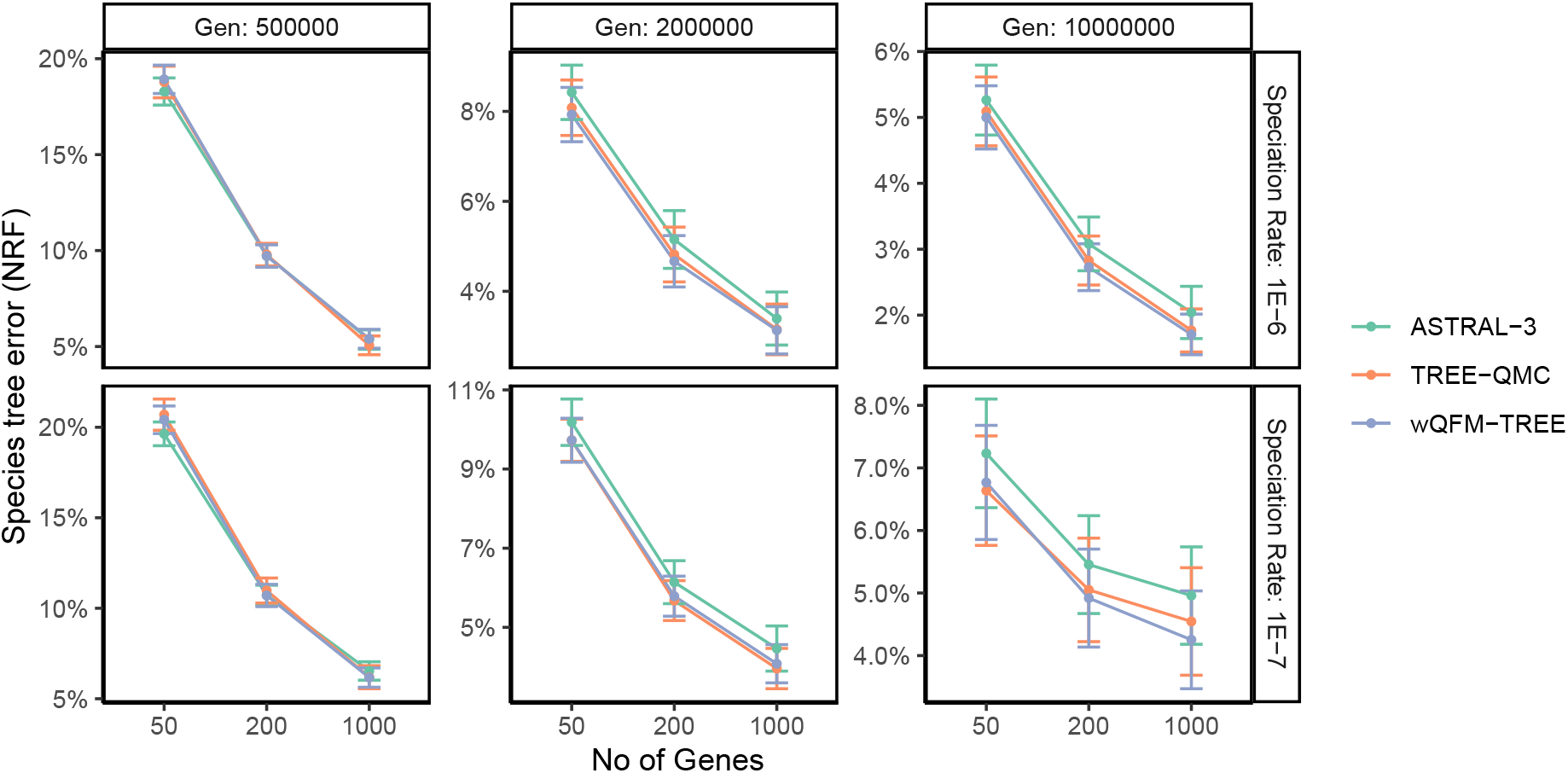
Comparison of methods on 200-taxon dataset with varying tree shapes and numbers of genes.

wQFM-TREE and TREE-QMC performed very closely on this particular dataset, with most of the differences being not statistically significant. Among the 18 model conditions examined, wQFM-TREE was better than TREE-QMC on 11 model conditions, with statistically significant differences in 2 of these 11 model conditions. On the other hand, TREE-QMC was better than wQFM-TREE on 6 model conditions, with improvements being statistically significant in 2 instances.

##### Results on varying numbers (200, 500, and 1000) of taxa

Figure 3 shows the assessment of different methods on varying the number of taxa (200, 500, and 1000) and genes (50, 200, and 1000) with the tree shape fixed to 2 M/1e-6. Remarkably, wQFM-TREE outperformed ASTRAL in all the model conditions, with differences being statistically significant (*P <* 0.05) in 8 of the 9 model conditions. Notably, the superiority of wQFM-TREE over ASTRAL became more pronounced with the increasing number of taxa. It is important to note that ASTRAL is a highly accurate and the most widely-used method for species tree estimation. Therefore, the demonstrated consistent improvement over ASTRAL, albeit small in some cases, is remarkable. wQFM-TREE and TREE-QMC showed comparable performance, with no statistically significant difference.

**Figure 3:**
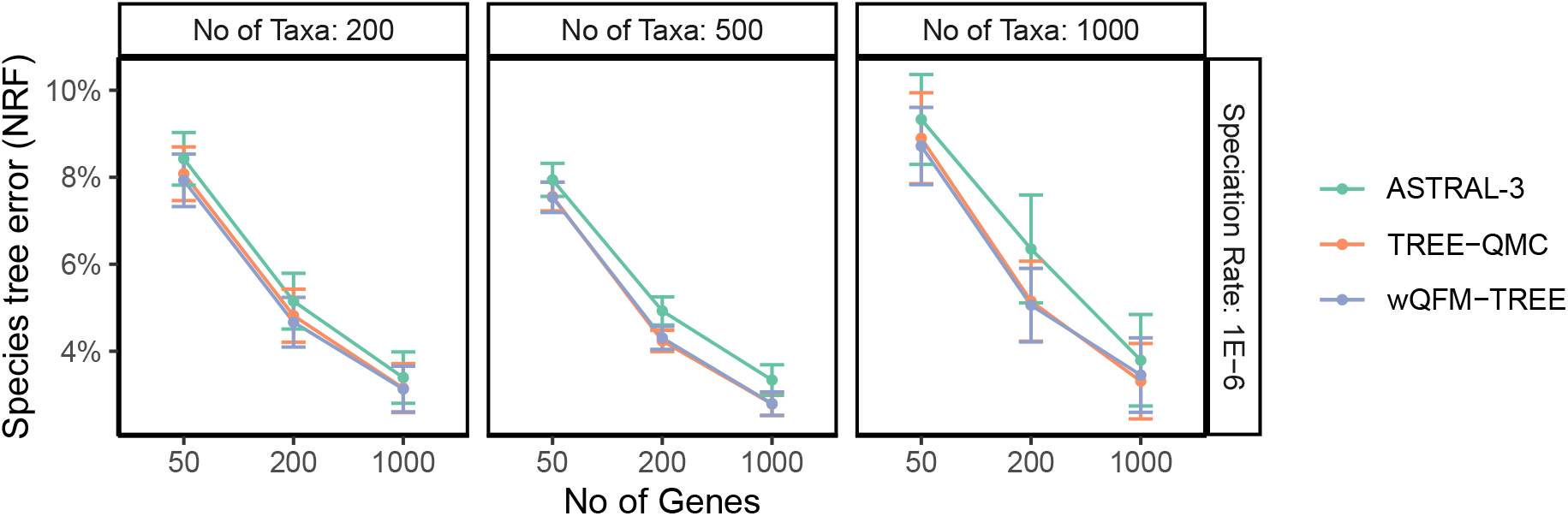
Comparison of methods on varying numbers of taxa (200, 500, and 1000) and genes (50, 200, and 1000) with the tree shape fixed to 2M/1e-6.

#### 4.1.3 Handling polytomies

wQFM-TREE can handle polytomies, making it suitable for partially resolved gene trees. Thus, wQFM-TREE can be applied even after removing low-support branches. However, similar to ASTRAL, the presence of polytomies increases the running time. In the presence of substantial amounts of polytomies, resolving them arbitrarily proves to be an effective strategy, resulting in a significant reduction in running time without sacrificing accuracy. As we have observed in our study, resolving the polytomies does not have any notable impact on the species tree accuracy (Supplementary Figure 4), while significantly improving the running time of wQFM-TREE (Supplementary Table 1).

### 4.2 Results on empirical dataset

We have re-analyzed the transcriptome dataset of 1178 species and 410 gene trees of plant species. The tree constructed by wQFM-TREE, shown in Figure 4, is highly congruent with the tree estimated by ASTRAL-III v.5.7.8 which was presented in [1kp, 2019]. wQFM-TREE successfully identified all the major clades, and the majority of the recovered relations between different clades are supported by other popular methods and do not violate any known consensus among scientists. The key relationships are discussed below.

**Figure 4:**
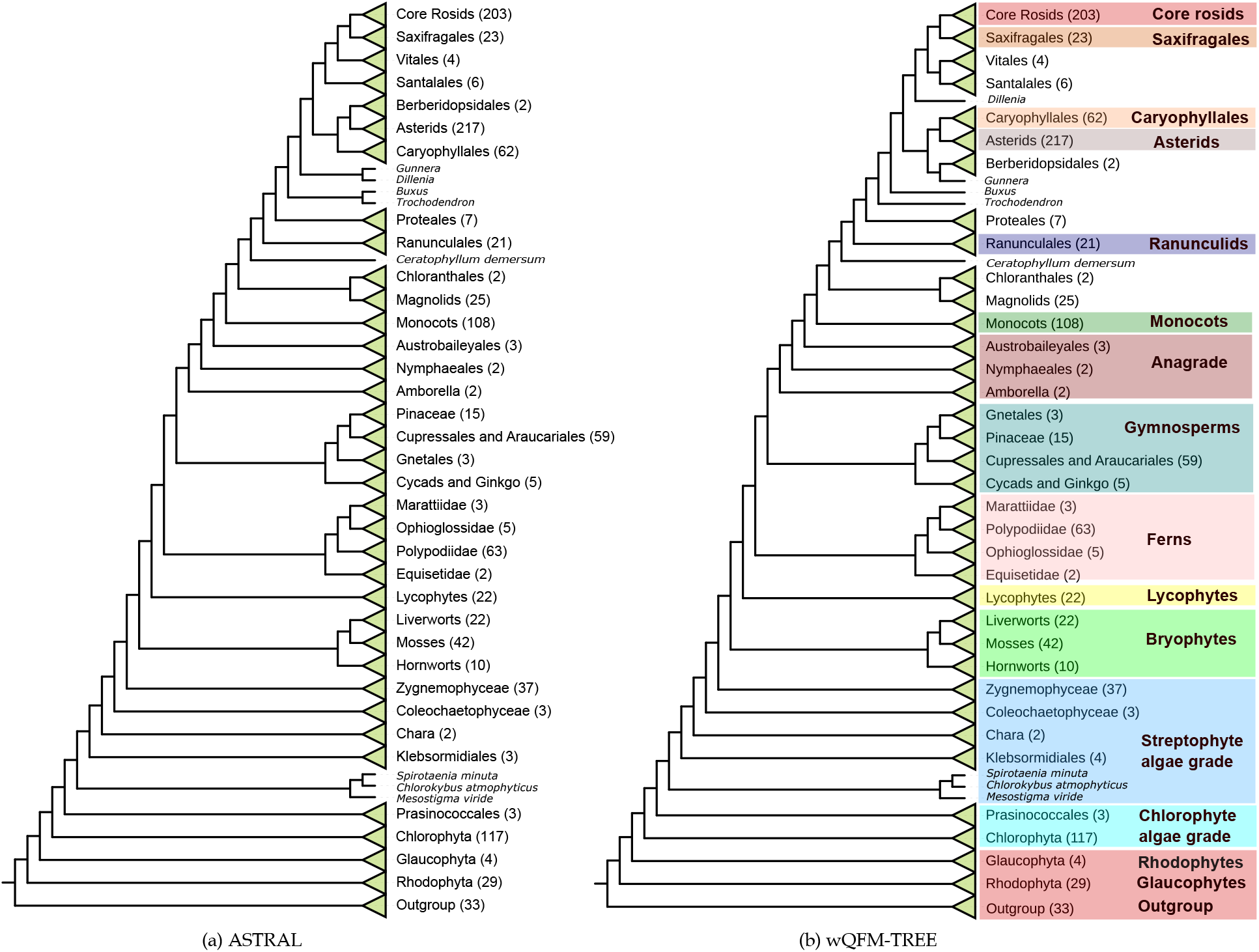
Phylogenetic trees reconstructed by wQFM-TREE using 410 single-copy nuclear gene families extracted from genome and transcriptome data from 1,178 species, including green plants (Viridiplantae), glaucophytes, and red algae. Species numbers are shown for each lineage.

#### Primary acquisition of plastid

Relationship among Viridiplantae, Glaucophyta, and Rhodo-phyta is crucial because it is related to the acquisition of plastid which is a pivotal event in the history of life. The sister relationship of Viridiplantae and Glaucophyta found here, indicating that ancestral red algae lost flagella and peptidoglycan biosynthesis, is aligned with the ASTRAL-tree presented in [1kp, 2019].

#### Viridiplantae

wQFM-TREE recovered the monophyletic relationship of Viridiplantae with early diverging Cholorophyta and Strepto-phyta which is consistent with prior studies. The placement of Prasinococcales is contentious and was found to be unstable in [1kp, 2019]. However, the placement of Prasinococcales by wQFM-TREE is identical to the ASTRAL-estimated tree.

#### Diversification within Chlorophyta and Streptophyta

wQFM-TREE successfully reconstructed the monophyletic relationships within Trebouxiophyceae, Chlorophyceae, and Ulvophyceae, except Briopsidales (an order of green algae, in the class Ulvophyceae). Briopsidales did not exhibit monophyly with other Ulvophyceae; instead, it was positioned as a sister group to Pedinophyceae with a very low support (26.8%). ASTRAL also did not place Briopsidales as a sister to other Ulvophyceae. Trebouxiophyceae was placed as a sister to a clade containing Chlorophyceae and other Ulvophyceae. In the case of Streptophyta, our analysis, identical to ASTRAL, recovered *Mesostigma, Spirotaenia minuta* and *Chlorokybus* in a clade that is sister to the remainder of Streptophyta with successive divergence of Klebsormidiales, Charophyceae, Coleochaetophyceae and Zygnematophyceae relative to Embryophyta.

#### Embryophyta

Although the relationships among the Bryophytes (mosses, liverworts, and hornworts) have been contentious and resolved as a grade in some studies, wQFM-TREE recovered Bryophytes as a monophyletic group similar to the ASTRAL-estimated tree and rejected the hypothesis that liverworts are sister to all other extant land plant lineages.

#### Vascular plants

Lycophytes were correctly recovered as the sister group of ferns and seed plants. There are conflicting results between supermatrix, plastom-based trees, and ASTRAL in the placement of Marattiales. wQFM-TREE supported the result of ASTRAL by placing it as a sister of Polypodiidae.

#### Seed plants

wQFM-TREE successfully recovered various groups/subgroups in seed plants. Gymnosperms were correctly recovered as sister to flowering plants. In the case of gymnosperms, the placement of Gnetales conflicts strongly among different methods, which gives rise to the ‘Gnecup’, ‘Gnepine’ and ‘Gnetifer’ hypotheses. wQFM-TREE supported the ‘Gnepine’ hypothesis, albeit with a low support of 41%. It placed Gnetales as sisters to Pinales which is in agreement with supermatrix analysis. However, it differed from ASTRAL, which strongly supports the Gnetifer hypotheses, i.e., Gnetales is sister to Conifers (Araucariales, Cupressales and Pinales) as a whole.

In the case of Angiosperms, Amborellales, Nymphaeales and Austrobaileyales were placed as successive sisters to all other angiosperms. Similar to the ASTRAL analyses, Chloranthales and Magnoliids were placed as sister groups. The placements of Core rosids, Saxifragales, and Asterids were also aligned with the ASTRAL-estimated tree.

### 4.3 Running time

We performed the experiments on a Linux machine with 64 GB RAM and Intel(R) Core(TM) i7-10700K 3.80GHz processor. The runtimes required by different methods on various datasets are presented in Table 1 in the Supplementary Materials.

The running times of wQFM-TREE, ASTRAL, and TREE-QMC suggest that all of them are fast enough to handle large datasets containing thousands of taxa and genes. While their run times are comparable for relatively small numbers of taxa, both ASTRAL and TREE-QMC are faster than wQFM-TREE on larger datasets. However, wQFM-TREE remains fast enough to analyze the largest dataset analyzed in this study with 2000 taxa and 1000 genes in about 6 hours. We report the running time of wQFM-TREE on fully-resolved gene trees, indicating that resolving polytomies substantially reduces runtime without sacrificing accuracy (Supplementary Figure 4).

## 5 Conclusions

In this work, we introduced wQFM-TREE, a novel method that allows the application of wQFM to gene trees without the need to decompose them into induced quartets. This innovation significantly enhances the scalability of wQFM-TREE, allowing it to handle datasets with thousands of taxa and genes efficiently. Our analyses, including the examination of the green plant dataset with over a thousand species, underscored the scalability of wQFM-TREE. Importantly, we have proposed several novel techniques, drawing from algorithms and combinatorics, for computing various quartet-based metrics (e.g., the number of satisfied or violated quartets) without explicitly enumerating the quartets in a given set of gene trees. These techniques not only enhance the scalability of wQFM but are also expected to facilitate quartet-based scoring techniques in general, eliminating the need to explicitly enumerate quartets in gene tree sets.

While ASTRAL has been a widely adopted choice, alternative heuristics like wQFM have faced challenges despite being more accurate than ASTRAL, as the running time becomes prohibitively large with increasing numbers of taxa. The techniques presented here address this scalability issue and make wQFM suitable to analyze very large datasets without sacrificing its accuracy. wQFM-TREE was consistently found to be more accurate or as good as ASTRAL and TREE-QMC. Therefore, we believe that wQFM-TREE and the underlying novel techniques represent a significant advancement in the field of quartet-based phylogeny estimation.

Although at this stage, TREE-QMC is considerably faster than both ASTRAL and wQFM-TREE on large datasets, there is considerable potential for algorithmic improvements of wQFM-Tree, such as consolidating the input gene trees into a unified data structure to avoid redundant calculations, employing heuristic strate-gies to reduce the number of FM iterations, etc. The partitioning technique of the QFM algorithm is inherently parallelizable, and future studies need to develop a parallel version of wQFM-TREE. The scalability of wQFM-TREE can be further enhanced by the divide-and-conquer framework proposed for making summary methods faster [Bayzid et al., 2014]. wQFM-TREE can handle single-copy rooted and unrooted gene trees. Extending it for multi-copy gene trees would be an interesting future research direction.

## Availability

wQFM-TREE is freely available in open source form at https://github.com/abdur-rafi/wQFM-TREE. All the datasets analyzed in this paper are from previously published studies and are publicly available.

