## Supplementary Material for "wQFM-TREE: highly accurate and scalable quartet-based species tree inference from gene trees"

These supplementary materials present additional details about the proposed technique for computing scores of candidate bipartitions from gene trees, the time complexity of wQFM-TREE, and additional results, figures, and tables.

### 1 Divide and conquer approach of wQFM

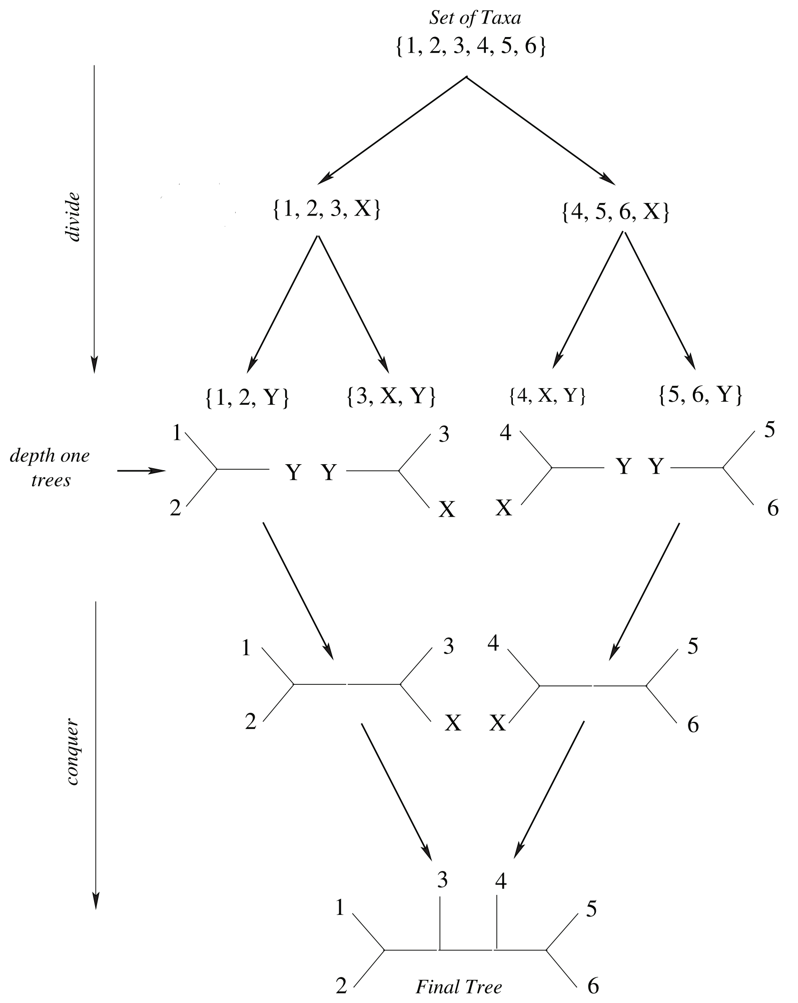

Figure 1: **Divide and conquer approach of wQFM**. Divide: At each step, the input set of taxa of this step is partitioned into two sets and a unique dummy taxon is added to both of the sets. These sets are inputs to the successive divide steps. If at any step, the size of the taxa set becomes less than or equal to 3, a depth one tree over the taxa set is returned. Conquer: At each step, there are two trees corresponding to the divide calls initiated at this step. These two trees are joined on the dummy taxon introduced at this step during divide. For example, the leftmost two depth one trees, when returned to its caller, are joined on the dummy taxon Y .

### 2 Additional details about score computation

**Lemma 1.** *$S^{(g,u)}$ for all internal nodes $u$ of gene tree $g$, form a partition of $S^{(g)}$, the set of satisfied resolved quartets of gene tree $g$.*

*Proof.* To prove the statement, we need to show that the union of $S^{(g,u)}$ over all internal nodes $u$ is $S^{(g)}$ and any pair of $S^{(g,u)}$ and $S^{(g,v)}$, for two distinct nodes $u$ and $v$, are disjoint.

The proof is quite simple. First, we show that the union of $S^{(g,u)}$ over all internal nodes $u$ is $S^{(g)}$. We consider a satisfied (resolved) quartet $ab|cd$ where $a, b \in F_A$; $c, d \in F_B$. Each quartet of leaves in the gene tree defines two anchor nodes. For the quartet $ab|cd$, one anchor node is directly connected to $a, b$ and the other to $c, d$ in the quartet tree. Let $z$ be the internal node, which is the anchor directly connected to $a, b$. Of course, $a$ and $b$ belong to two distinct components in $C^{(g,z)}$, and both $c, d$ pertain to a third component of $C^{(g,z)}$. Thus, every quartet belongs to $S^{(g,u)}$ for at least one internal node $u$. Thus, the union of $S^{(g,u)}$ over all internal nodes $u$ is $S^{(g)}$.

Now, we show that any pair of $S^{(g,u)}$ and $S^{(g,v)}$, for two distinct nodes $u$ and $v$, are disjoint. Again, we consider a satisfied (resolved) quartet $ab|cd$ where $a, b \in F_A$; $c, d \in F_B$ and let $z$ be the internal node, which is the anchor directly connected to $a, b$. We consider the path from $a$ to $b$ in gene tree $g$. For any internal node $x$ outside of the path, $a$ and $b$ are in the same component element of $C^{(g,x)}$, which violates the definition of $S^{(g,x)}$. For any node $x$ in the path except $z$, nodes $a, c, d$ or $b, c, d$ will be in the same component element of $C^{(g,x)}$. Therefore, the quartet $ab|cd$ belongs to only $S^{(g,z)}$. Therefore, a quartet $ab|cd$ can only belong to $S^{(g,u)}$ for one internal node $u$ which is the anchor node defined in the gene tree by the leaves $a, b, c, d$ and directly connected to $a, b$ in the quartet tree. Thus, any pair of $S^{(g,u)}$ and $S^{(g,v)}$, for two different nodes $u$ and $v$, are disjoint. This completes our proof.

□

#### 2.1 Additional details about weight computation

Let $F_A^{(g,u,i)}$ be the set of real taxa of $F_A$ that are in $C_i^{(g,u)}$, $X_R^{(g,u,i)}$ be the set of real taxa in $C_i^{(g,u)}$ that are under a dummy taxon $X$ and $R_A^{(g,u,i)}$ be the set of real taxa of $R_A$ that are in $C_i^{(g,u)}$. $F_B^{(g,u,i)}$ and $R_A^{(g,u,i)}$ are similarly defined.

##### 2.1.1 Computing $w(PA_{i,j}^{(g,u)})$ and $w(PB_{i,j}^{(g,u)})$

Let $P_1 = \{\{a, b\} : a \in F_A^{(g,u,i)}, b \in F_A^{(g,u,j)}\}$. $P_1$ contains all elements of $PA_{i,j}^{(g,u)}$. However, it also contains the unordered pairs $\{a, b\}$ where both $a$ and $b$ are under the same dummy taxon. Let $P_2 = \bigcup_{X \in D_A} \{\{a, b\} : a \in X_R^{(g,u,i)}, b \in X_R^{(g,u,j)}\}$. $P_2$ contains all the pairs $\{a, b\}$ where $a \in F_A^{(g,u,i)}$, $b \in F_A^{(g,u,j)}$ and $a, b$ both are under the same dummy taxon. Now, $P_2$ is a subset of $P_1$ and $PA_{i,j}^{(g,u)} = P \backslash Q$. Thus, $w(PA_{i,j}^{(g,u)}) = w(P_1) - w(P_2)$.

Now $w(P_1) = w(F_A^{(g,u,i)}) w(F_A^{(g,u,j)})$ and $w(P_2) = \sum_{X \in D_A} w(X_R^{(g,u,i)}) w(X_R^{(g,u,j)})$. Thus, we get the formula of $w(PA_{i,j}^{(g,u)})$ as follows.

$$w(PA_{i,j}^{(g,u)}) = w(F_A^{(g,u,i)})w(F_A^{(g,u,j)}) - \sum_{X \in D_A} w(X_R^{(g,u,i)})w(X_R^{(g,u,j)})$$

We can compute $w(F_A^{(g,u,i)})$ and $w(X_R^{(g,u,i)})$ for all internal nodes $u$ of gene tree $g$ through a single post-order traversal of $g$ after rooting it arbitrarily.

Similarly,

$$w(PB_{i,j}^{(g,u)}) = w(F_B^{(g,u,i)})w(F_B^{(g,u,j)}) - \sum_{X \in D_B} w(X_R^{(g,u,i)})w(X_R^{(g,u,j)})$$

##### 2.1.2 Computing $w(PB_k^{(g,u)})$

To compute $w(PB_k^{(g,u)})$, we can follow a similar approach as we did for $w(PB_{i,j}^{(g,u)})$, with some minor modifications.

$$\begin{aligned} w(PB_k^{(g,u)}) &= \frac{1}{2}(w(F_B^{(g,u,k)})w(F_B^{(g,u,k)}) - w(R_B^{(g,u,k)}) - \sum_{X \in D_B} w(X_R^{(g,u,k)})w(X_R^{(g,u,k)})) \\ &= \frac{1}{2}(w(F_B^{(g,u,k)})^2 - w(R_B^{(g,u,k)}) - \sum_{X \in D_B} w(X_R^{(g,u,k)})^2) \end{aligned}$$

This expression differs from that of $w(PB_{i,j}^{(g,u)})$ for a few reasons. Unlike the case of $w(PB_{i,j}^{(g,u)})$, $w(F_B^{(g,u,k)})^2$ also contains the sum of weights of $\{a, a\}, a \in F_B^{(g,u,k)}$. We do not include $\{a, a\}$ by definition in $PB_k^{(g,u)}$ as a real taxon cannot participate twice in a quartet. Weights of $\{a, a\}$ where $a \in X_R$ for some $X \in D_B$ are already included in $\sum_{X \in D_B} w(X_R^{(g,u,k)})^2$. Therefore, additionally, we need to subtract weights of $\{a, a\}$ where $a$ is not under any dummy taxon, that is, $a \in R_B^{(g,u,k)}$. As $w(a) = 1$ for $a \in R_B^{(g,u,k)}$, sum of weights of these $\{a, a\}$ is given by $w(R_B^{(g,u,k)})$. Moreover, since both taxa are from the same set, the weight of all unordered pairs is considered twice. Hence, by dividing the expression by 2, we obtain the final formula.

##### 2.1.3 Efficient formulation of $w(S^{(g,u)})$

We compute $w(S^{(g,u)})$ efficiently in the following way. Let $w(PB^{(g,u)}) = \sum_k w(PB_k^{(g,u)})$. For a certain $i, j$,

$$\begin{aligned} \sum_{k \neq i,j} w(S_{i,j,k}^{(g,u)}) &= \sum_{k \neq i,j} w(PA_{i,j}^{(g,u)})w(PB_k^{(g,u)}) \\ &= w(PA_{i,j}^{(g,u)}) \sum_{k \neq i,j} w(PB_k^{(g,u)}) \\ &= w(PA_{i,j}^{(g,u)}) \cdot (w(PB^{(g,u)}) - w(PB_i^{(g,u)}) - w(PB_j^{(g,u)})) \end{aligned}$$

Thus, we get

$$
\begin{aligned}
w(S^{(g,u)}) &= \sum_{i<j} \sum_{k \neq i,j} w(S^{(g,u)}_{i,j,k}) \\
&= \sum_{i<j} w(PA^{(g,u)}_{i,j}) \cdot (w(PB^{(g,u)}) - w(PB^{(g,u)}_{i}) - w(PB^{(g,u)}_{j}))
\end{aligned}
$$

##### 2.1.4 Computing $w(PA^{(g)})$ and $w(PB^{(g)})$

Similar to 2.1.2, $PA^{(g)}$ contains pairs $\{a, b\}$ where both $a, b$ are from the same set $\mathcal{X}^{(g)} \cap F_A$. Therefore,

$$
w(PA^{(g)}) = \frac{1}{2}(w(F_A \cap \mathcal{X}^{(g)})^2 - w(R_A \cap \mathcal{X}^{(g)}) - \sum_{X \in D_A} w(X_R \cap \mathcal{X}^{(g)})^2)
$$

Similarly,

$$
w(PB^{(g)}) = \frac{1}{2}(w(F_B \cap \mathcal{X}^{(g)})^2 - w(R_B \cap \mathcal{X}^{(g)}) - \sum_{X \in D_B} w(X_R \cap \mathcal{X}^{(g)})^2)
$$

##### 2.1.5 Efficient formulation of $w(U^{(g,u)})$

$$
w(U^{(g,u)}) = \sum_{i<j} w(PA^{(g,u)}_{i,j}) ( \sum_{k<l; k,l \notin \{i,j\}} w(PB^{(g,u)}_{k,l}))
$$

Let $SB^{(g,u)}_i = \sum_{k<l;(k=i)\vee(l=i)} w(PB^{(g,u)}_{k,l})$ and $SB^{(g,u)} = \sum_{k<l} w(PB^{(g,u)}_{k,l})$. Utilizing these definitions, we can refine the expression as follows.

$$
\sum_{k<l; k,l \notin \{i,j\}} w(PB^{(g,u)}_{k,l}) = SB^{(g,u)} - SB^{(g,u)}_i - SB^{(g,u)}_j + w(PB^{(g,u)}_{i,j})
$$

Thus, we obtain the following equation.

$$
\begin{aligned}
w(U^{(g,u)}) &= \sum_{i<j} w(PA^{(g,u)}_{i,j}) ( \sum_{k<l; k,l \notin \{i,j\}} w(PB^{(g,u)}_{k,l})) \\
&= \sum_{i<j} w(PA^{(g,u)}_{i,j}) (SB^{(g,u)} - SB^{(g,u)}_i - SB^{(g,u)}_j + w(PB^{(g,u)}_{i,j}))
\end{aligned}
$$

#### 2.2 Example demonstrating the score computation method

We demonstrate the score calculation process by calculating the score of the bipartition $(A, B) = (\{1, 4, 12, X\}, \{2, 6, 7, Y\})$ for the unrooted gene tree $g$ shown in Figure 2a. Here, $X$ and $Y$ are dummy taxa. Their tree structures are shown in Figure 2a. From the tree structure, $X_R = \{5, 10, 11\}$ and $Y_R = \{3, 8, 9\}$. The weights of all the real taxa which are not under X or Y (taxon 1, 2, 4, 6, 7, 12) are 1. Using the weighting mechanism described in Section 2.3 of the main paper, we calculate the weights of the taxa in $X_R$ and $Y_R$.

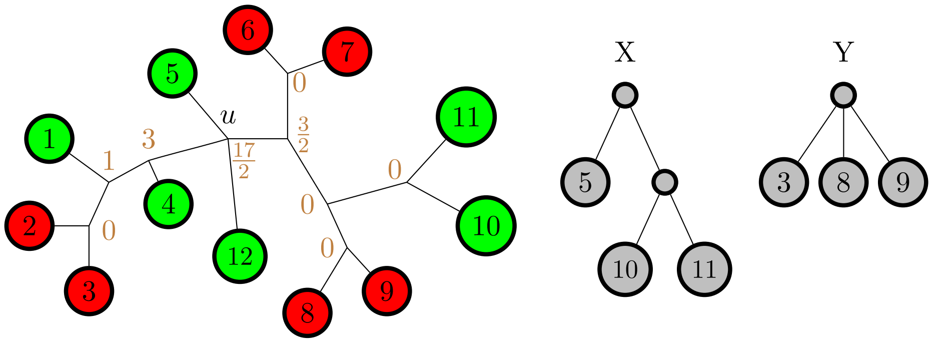

(a) Example gene tree and the tree structure of the dummy taxa $X$ and $Y$ . The values at each internal node of the gene tree denote the value of $w(S^{(g,u)})$ for that node.

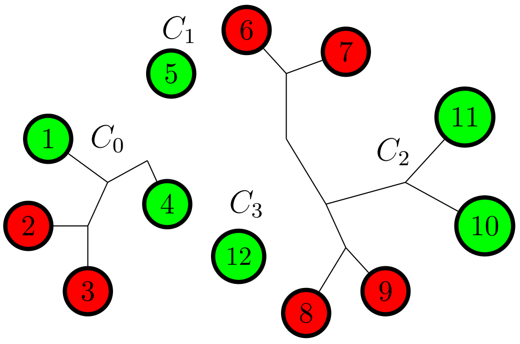

(b) Resulting components after removing the internal node $u$. The components are labeled $C_0 - C_3$ in an arbitrary order.

Figure 2: An example gene tree to demonstrate our score calculation process for the bipartition $(\{1, 4, 12, X\}, \{2, 6, 7, Y\})$.

For taxa in $X_R$, $w(5) = \frac{1}{2}$, $w(10) = \frac{1}{4}$, and $w(11) = \frac{1}{4}$. For taxa in $Y_R$, $w(3) = \frac{1}{3}$, $w(8) = \frac{1}{3}$, and $w(9) = \frac{1}{3}$.

From $A$ and $B$, $R_A = \{1, 4, 12\}$,$D_A = \{X\}$, $F_A = \{1, 4, 5, 10, 11, 12\}$, $R_B = \{2, 6, 7\}$,$D_B = \{Y\}$, $F_B = \{2, 3, 6, 7, 8, 9\}$.

We recall that,

$$Score(A, B, \mathcal{G}) = \sum_{g \in \mathcal{G}} (2w(S^{(g)}) - w(S^{(g)} \cup V^{(g)} \cup U^{(g)}) + w(U^{(g)}))$$

Here, $\mathcal{G}$ is a set of unrooted gene trees. We have one gene tree $g$ in $\mathcal{G}$ in our example. Therefore, the equation becomes,

$$Score(A, B, \mathcal{G}) = 2w(S^{(g)}) - w(S^{(g)} \cup V^{(g)} \cup U^{(g)}) + w(U^{(g)})$$

We show the detailed calculation of $w(S^{(g,u)})$ and $w(U^{(g,u)})$ only for the internal node $u$, where $u$ is the marked node in Figure 2a. Removing $u$ creates 4 components, $C_0^{(g,u)}$, $C_1^{(g,u)}$, $C_2^{(g,u)}$, $C_3^{(g,u)}$. These components are shown in Figure 2b, labeled by $C_0, C_1, C_2, C_3$.

##### 2.2.1 Computing $w(S^{(g)})$

We determine $w(S^{(g)}) = \sum_{v \in g} w(S^{(g,v)})$. We present the detailed computation of $w(S^{(g,u)})$. We recall that,

$$\begin{aligned} w(S^{(g,u)}) &= \sum_{i<j} \sum_{k \neq i,j} w(S_{i,j,k}^{(g,u)}) \\ &= \sum_{i<j} w(PA_{i,j}^{(g,u)}) \cdot (w(PB^{(g,u)}) - w(PB_i^{(g,u)}) - w(PB_j^{(g,u)})) \end{aligned}$$

To compute $w(S^{(g,u)})$, we need to determine $w(PA_{i,j}^{(g,u)})$ for each $i < j$ and $w(PB_i^{(g,u)})$ for each $i$ . For this task, We calculate the value of all the required expressions as shown below.

$F_A^{(g,u,0)} = \{1, 4\}$, $F_A^{(g,u,1)} = \{5\}$, $F_A^{(g,u,2)} = \{10, 11\}$, $F_A^{(g,u,3)} = \{12\}$.
$w(F_A^{(g,u,0)}) = 2$, $w(F_A^{(g,u,1)}) = \frac{1}{2}$, $w(F_A^{(g,u,2)}) = \frac{1}{2}$, $w(F_A^{(g,u,3)}) = 1$.
$F_B^{(g,u,0)} = \{2, 3\}$, $F_B^{(g,u,1)} = \{\}$, $F_B^{(g,u,2)} = \{6, 7, 8, 9\}$, $F_B^{(g,u,3)} = \{\}$.
$w(F_B^{(g,u,0)}) = \frac{4}{3}$, $w(F_B^{(g,u,1)}) = 0$, $w(F_B^{(g,u,2)}) = \frac{8}{3}$, $w(F_B^{(g,u,3)}) = 0$.
$R_A^{(g,u,0)} = \{1, 4\}$, $R_A^{(g,u,1)} = \{\}$, $R_A^{(g,u,2)} = \{\}$, $R_A^{(g,u,3)} = \{12\}$.
$w(R_A^{(g,u,0)}) = 2$, $w(R_A^{(g,u,1)}) = 0$, $w(R_A^{(g,u,2)}) = 0$, $w(R_A^{(g,u,3)}) = 1$.
$R_B^{(g,u,0)} = \{2\}$, $R_B^{(g,u,1)} = \{\}$, $R_B^{(g,u,2)} = \{6, 7\}$, $R_B^{(g,u,3)} = \{\}$
$w(R_B^{(g,u,0)}) = 1$, $w(R_B^{(g,u,1)}) = 0$, $w(R_B^{(g,u,2)}) = 2$, $w(R_B^{(g,u,3)}) = 0$.
$X_R^{(g,u,0)} = \{\}$, $X_R^{(g,u,1)} = \{5\}$, $X_R^{(g,u,2)} = \{10, 11\}$, $X_R^{(g,u,3)} = \{\}$.
$w(X_R^{(g,u,0)}) = 0$, $w(X_R^{(g,u,1)}) = \frac{1}{2}$, $w(X_R^{(g,u,2)}) = \frac{1}{2}$, $w(X_R^{(g,u,3)}) = 0$.
$Y_R^{(g,u,0)} = \{3\}$, $Y_R^{(g,u,1)} = \{\}$, $Y_R^{(g,u,2)} = \{8, 9\}$, $Y_R^{(g,u,3)} = \{\}$.
$w(Y_R^{(g,u,0)}) = \frac{1}{3}$, $w(Y_R^{(g,u,1)}) = 0$, $w(Y_R^{(g,u,2)}) = \frac{2}{3}$, $w(Y_R^{(g,u,3)}) = 0$.

Now, we can compute $w(PA_{i,j}^{(g,u)})$ for each $i < j$

$$w(PA_{0,1}^{(g,u)}) = w(F_A^{(g,u,0)}) w(F_A^{(g,u,1)}) - w(X_R^{(g,u,0)}) w(X_R^{(g,u,1)}) = 2 \cdot \frac{1}{2} - 0 \cdot \frac{1}{2} = 1.$$

Similarly,

$$w(PA^{(g,u)}_{0,2}) = 1, w(PA^{(g,u)}_{0,3}) = 2, w(PA^{(g,u)}_{1,2}) = 0, w(PA^{(g,u)}_{1,3}) = \frac{1}{2}, w(PA^{(g,u)}_{2,3}) = \frac{1}{2}.$$

Then, we compute $w(PB^{(g,u)}_k)$ for each $k$.

$$w(PB^{(g,u)}_0) = \frac{1}{2}(w(F^{(g,u,0)}_B)^2 - w(R^{(g,u,0)}_B) - w(Y^{(g,u,0)}_R)^2) = \frac{1}{2}((\frac{4}{3})^2 - 1 - \frac{1}{3^2}) = \frac{1}{3}.$$

Similarly,

$$w(PB^{(g,u)}_1) = 0, w(PB^{(g,u)}_2) = \frac{7}{3}, w(PB^{(g,u)}_3) = 0.$$

Then, we compute the value of $PB^{(g,u)}$.

$$PB^{(g,u)} = w(PB^{(g,u)}_0) + w(PB^{(g,u)}_1) + w(PB^{(g,u)}_2) + w(PB^{(g,u)}_3) = \frac{8}{3}$$

We are done computing all the necessary values required to compute $\sum_{k\neq i,j} w(S^{(g,u)}_{i,j,k})$ for a fixed $i, j$.

$$\sum_{k\neq i,j} w(S^{(g,u)}_{i,j,k}) = w(PA^{(g,u)}_{i,j}) \cdot (w(PB^{(g,u)}) - w(PB^{(g,u)}_i) - w(PB^{(g,u)}_j))$$

$$\begin{aligned}\sum_{k\neq 0,1} w(S^{(g,u)}_{0,1,k}) &= w(PA^{(g,u)}_{0,1}) \cdot (w(PB^{(g,u)}) - w(PB^{(g,u)}_0) - w(PB^{(g,u)}_1)) \\ &= 1 \cdot (\frac{8}{3} - \frac{1}{3} - 0) = \frac{7}{3}\end{aligned}$$

Similarly,

$$\sum_{k\neq 0,2} w(S^{(g,u)}_{0,2,k}) = 0, \sum_{k\neq 0,3} w(S^{(g,u)}_{0,3,k}) = \frac{14}{3}, \sum_{k\neq 1,2} w(S^{(g,u)}_{1,2,k}) = 0,$$
$$\sum_{k\neq 1,3} w(S^{(g,u)}_{1,3,k}) = \frac{4}{3}, \sum_{k\neq 2,3} w(S^{(g,u)}_{2,3,k}) = \frac{1}{6},$$

Finally, we compute $w(S^{(g,u)})$.

$$\begin{aligned}w(S^{(g,u)}) =& \sum_{i<j} \sum_{k\neq i,j} S^{(g,u)}_{i,j,k} \\ =& \sum_{k\neq 0,1} w(S^{(g,u)}_{0,2,k}) + \sum_{k\neq 0,2} w(S^{(g,u)}_{0,2,k}) + \sum_{k\neq 0,3} w(S^{(g,u)}_{0,3,k}) + \sum_{k\neq 1,2} w(S^{(g,u)}_{1,2,k}) + \\ & \sum_{k\neq 1,3} w(S^{(g,u)}_{1,3,k}) + \sum_{k\neq 2,3} w(S^{(g,u)}_{2,3,k}) \\ =& \frac{7}{3} + 0 + \frac{14}{3} + 0 + \frac{4}{3} + \frac{1}{6} = \frac{17}{2}\end{aligned}$$

By performing similar calculations for the other nodes, we obtain the value of $w(S^{(g)})$.

$$\begin{aligned}w(S^{(g)}) &= \sum_{u\in g} w(S^{(g,u)}) \\ &= 0 + 1 + 3 + \frac{17}{2} + 0 + \frac{3}{2} + 0 + 0 + 0 = 14\end{aligned}$$

##### 2.2.2 Computing $w(S^{(g)} \cup V^{(g)} \cup U^{(g)})$

Now, we compute $w(S^{(g)} \cup V^{(g)} \cup U^{(g)})$. We need to compute $w(PA^{(g)})$ and $w(PA^{(g)})$ for this task.

Since we have no missing taxon, $\mathcal{X}^{(g)} = \mathcal{X}$, $F_A \cap \mathcal{X}^{(g)} = F_A$, $F_B \cap \mathcal{X}^{(g)} = F_B$, $R_A \cap \mathcal{X}^{(g)} = R_A$, $R_B \cap \mathcal{X}^{(g)} = R_B$, $X_R \cap \mathcal{X}^{(g)} = X_R$, and $Y_R \cap \mathcal{X}^{(g)} = Y_R$.

$$w(F_A \cap \mathcal{X}^{(g)}) = 4, w(F_B \cap \mathcal{X}^{(g)}) = 4, w(R_A \cap \mathcal{X}^{(g)}) = 3$$
$$w(R_B \cap \mathcal{X}^{(g)}) = 3, w(X_R \cap \mathcal{X}^{(g)}) = 1, w(Y_R \cap \mathcal{X}^{(g)}) = 1$$

$$\begin{aligned} w(PA^{(g)}) &= \frac{1}{2}(w(F_A \cap \mathcal{X}^{(g)})^2 - w(R_A \cap \mathcal{X}^{(g)}) - w(X_R \cap \mathcal{X}^{(g)})^2) \\ &= \frac{1}{2}(16 - 3 - 1) = 6 \end{aligned}$$

Similarly,

$$w(PB^{(g)}) = 6$$

Finally, we compute $w(S^{(g)} \cup V^{(g)} \cup U^{(g)})$

$$w(S^{(g)} \cup V^{(g)} \cup U^{(g)}) = w(PA^{(g)}) \cdot w(PB^{(g)}) = 6 \cdot 6 = 36$$

##### 2.2.3 Computing $w(U^{(g)})$

The only task remaining is to compute $w(U^{(g)})$. As $u$ is the only polytomy node, $w(U^{(g)}) = w(U^{(g,u)})$. Therefore, we only need to compute $w(U^{(g,u)})$. For this task, we compute $w(PB_{i,j}^{(g,u)})$ for all $i < j$.

$$w(PB_{0,1}^{(g,u)}) = w(F_B^{(g,u,0)})w(F_B^{(g,u,1)}) - w(Y_R^{(g,u,0)})w(Y_R^{(g,u,1)}) = 2 \cdot 0 - \frac{1}{3} \cdot 0 = 0.$$

Similarly,

$$\text{w(PB}_{0,2}^{(g,u)}) = \tfrac{10}{3}, w(PB_{0,3}^{(g,u)}) = 0, w(PB_{1,2}^{(g,u)}) = 0, w(PB_{1,3}^{(g,u)}) = 0, w(PB_{2,3}^{(g,u)}) = 0.$$

We recall that,

$$\begin{aligned} w(U^{(g,u)}) &= \sum_{i<j} \sum_{k<l; k,l \notin \{i,j\}} w(U_{i,j,k,l}^{(g,u)}) \\ &= \sum_{i<j} \sum_{k<l; k,l \notin \{i,j\}} w(PA_{i,j}^{(g,u)}) w(PB_{k,l}^{(g,u)}) \\ &= \sum_{i<j} w(PA_{i,j}^{(g,u)}) \Big( \sum_{k<l; k,l \notin \{i,j\}} w(PB_{k,l}^{(g,u)}) \Big) \\ &= \sum_{i<j} w(PA_{i,j}^{(g,u)}) (SB^{(g,u)} - SB_i^{(g,u)} - SB_j^{(g,u)} + w(PB_{i,j}^{(g,u)})) \end{aligned}$$

We determine $SB^{(g,u)}$ and $SB_i^{(g,u)}$ for all $i$.

$$\begin{aligned}SB^{(g,u)} &= \sum_{k<l} w(PB_{k,l}^{(g,u)}) \\ &= w(PB_{0,1}^{(g,u)}) + w(PB_{0,2}^{(g,u)}) + w(PB_{0,3}^{(g,u)}) + w(PB_{1,2}^{(g,u)}) + w(PB_{1,3}^{(g,u)}) + w(PB_{2,3}^{(g,u)}) \\ &= 0 + \frac{10}{3} + 0 + 0 + 0 + 0 = \frac{10}{3}\end{aligned}$$

$$\begin{aligned}SB_0^{(g,u)} &= \sum_{k<l;(k=0)\vee(l=0)} w(PB_{k,l}^{(g,u)}) \\ &= w(PB_{0,1}^{(g,u)}) + w(PB_{0,2}^{(g,u)}) + w(PB_{0,3}^{(g,u)}) = 0 + \frac{10}{3} + 0 = \frac{10}{3}\end{aligned}$$

Similarly,

$$SB_1^{(g,u)} = 0, SB_2^{(g,u)} = \frac{10}{3}, SB_3^{(g,u)} = 0$$

With these values, we compute $\sum_{k,l\notin\{i,j\}} w(U_{0,1,i,j}^{(g,u)})$ for each $i < j$.

$$\begin{aligned}\sum_{k,l\notin\{0,1\}} w(U_{0,1,k,l}^{(g,u)}) &= w(PA_{0,1}^{(g,u)})(SB^{(g,u)} - SB_0^{(g,u)} - SB_1^{(g,u)} + w(PB_{0,1}^{(g,u)})) \\ &= 1 \cdot (\frac{10}{3} - \frac{10}{3} - 0 + 0) = 0\end{aligned}$$

Similarly,

$$\begin{aligned}&\sum_{k,l\notin\{0,2\}} w(U_{0,2,k,l}^{(g,u)}) = 0, \sum_{k,l\notin\{0,3\}} w(U_{0,3,k,l}^{(g,u)}) = 0, \sum_{k,l\notin\{1,2\}} w(U_{1,2,k,l}^{(g,u)}) = 0 \\ &\sum_{k,l\notin\{1,3\}} w(U_{1,3,k,l}^{(g,u)}) = \frac{5}{3}, \sum_{k,l\notin\{2,3\}} w(U_{2,3,k,l}^{(g,u)}) = 0\end{aligned}$$

Finally, we compute $w(U^{(g,u)})$.

$$\begin{aligned}w(U^{(g,u)}) = &\sum_{k,l\notin\{0,1\}} w(U_{0,1,k,l}^{(g,u)}) + \sum_{k,l\notin\{0,2\}} w(U_{0,2,k,l}^{(g,u)}) + \sum_{k,l\notin\{0,3\}} w(U_{0,3,k,l}^{(g,u)}) + \\ &\sum_{k,l\notin\{1,2\}} w(U_{1,2,k,l}^{(g,u)}) + \sum_{k,l\notin\{1,3\}} w(U_{1,3,k,l}^{(g,u)}) + \sum_{k,l\notin\{2,3\}} w(U_{2,3,k,l}^{(g,u)}) \\ =&0 + 0 + 0 + 0 + \frac{5}{3} + 0 = \frac{5}{3}\end{aligned}$$

As $u$ is the only internal node with polytomy,

$$w(U^{(g)}) = w(U^{(g,u)}) = \frac{5}{3}$$

Thus, our final score for the bipartition is,

$$Score(A, B, \mathcal{G}) = 2w(S^{(g)}) - w(S^{(g)} \cup V^{(g)} \cup U^{(g)}) + w(U^{(g)}) = 2 \cdot 14 - 36 + \frac{5}{3} = -\frac{19}{3}$$

#### 2.3 Weight normalization

Suppose a dummy taxon $X$ has $k$ real taxon under it. As gene trees contain only real taxa, we consider $X$ to be involved in a quartet if any of the real taxa under it participates in that quartet. As a result, $X$ can form $k$ times as many quartets as a real taxon $a \in R_A \cup R_B$ with the real taxa in a set $R \subseteq \mathcal{X}$ (assuming $a \notin R$, $X_R \cap R = \emptyset$ ). Since our scoring is based on sum of weights of quartets (described in Section 2.5 of the main paper), a dummy taxon gets prioritized in maximizing the score if no normalization is done (that is, all real taxa are assigned unit weight). Therefore, we attempt to ensure that a real taxon and a dummy taxon contribute equally to the score calculation.

For weight normalization, we can follow two approaches. One is to simply assign equal weights to all the real taxa under a dummy taxon. That is, if a dummy taxon $d$ has $k$ real taxa under it, then all of them are assigned weight $\frac{1}{k}$. Another approach is to assign weights according to the "dummy taxon tree structure" described in Section 2.3 of the main paper. Similar weighting mechanisms (uniform and non-uniform normalizations) were proposed and used in TREE-QMC [Han and Molloy, 2023]. The latter approach (i.e., non-uniform normalization) produced superior results in all our experiments and thus further supports the observations made in the TREE-QMC study.

#### 2.4 Gain calculation

We recall that wQFM algorithm improves the bipartition iteratively through transferring taxa from one partition to the opposite. The details of the procedure is described in [Reaz et al., 2014]. For a taxon $a \in A \cup B$, the corresponding gain, $G_a$, is the change in the score if $a$ is transferred to the opposite partition. That is, if $a \in A$, $G_a = \mathit{Score}(A\backslash\{a\}, B \cup \{a\}, \mathcal{G}) - \mathit{Score}(A, B, \mathcal{G})$. Our algorithm requires that we calculate gains of all the taxa present in $A$ and $B$. To do so, we calculate the change of $w(S^{(g,u)})$ at each internal node $u$ for each taxon $a \in A \cup B$ if it is transferred to the opposite partition. We also calculate the change of $w(U^{(g,u)})$ at each polytomy node $u$ and the change of $w(S^{(g)} \cup V^{(g)} \cup U^{(g)})$. From these, we determine the gain for each taxon.

We can be more efficient when dealing with gains of real taxa. We utilize the fact that the change in $w(S^{(g,u)})$ is the same for transferring any $a \in R_A$ within a component $C_i^{(g,u)}$ because all such $a$ has unit weight. Thus, we visit each internal node $u$, calculate and store the change of $w(S^{(g,u)})$ for each of its components. Then, these changes can be accumulated using a linear graph traversal to compute $G_a$ for all $a \in R_A$. Similarly, we can compute $G_a$ for all $a \in R_B$.

### 3 Time complexity

Let $n$ be the number of taxa present in the gene trees, $r$ and $d$ be the number of real taxa and dummy taxa present in a subproblem respectively. Let $(A, B)$ be the initial bipartition for the subproblem. Then, $r = |R_A| + |R_B|$ and $d = |D_A| + |D_B|$. We assume that the degree of the internal nodes is bounded by a constant $c$.

First, we can compute $w(F_A^{(g,u,i)})$, $w(F_B^{(g,u,i)})$ and $w(X_R^{(g,u,i)})$ for all $u$ in $O(nd)$ for a tree $g$. Therefore, for $k$ gene trees, the time complexity is $O(knd)$.

Then, for an internal node $u$ of a gene tree $g$, we can compute $w(PA_{i,j}^{(g,u)})$, $w(PB_{i,j}^{(g,u)})$, $w(PB_k^{(g,u)})$, $\sum_{X \in D_A} w(X_R^{(g,u,i)})w(X_R^{(g,u,j)})$ and $\sum_{X \in D_B} w(X_R^{(g,u,i)})w(X_R^{(g,u,j)})$ for $1 \leq i,j,k \leq \deg(u)$ in $O(\deg(u)^2 d)$. Thus, for the whole set of gene trees, we can perform this computation in $O(c^2dnk)$ time. After that, we can calculate $w(S^{(g,u)})$ in $O(deg(u))$. Then, $\sum_{g \in \mathcal{G}} w(S^{(g,u)})$ can be computed in $O(cnk)$.

Now, we can calculate $w(S^{(g)} \cup V^{(g)} \cup U^{(g)})$ from $\mathcal{X}^{(g)}$ in $O(n)$ complexity. Overall, it takes $O(nk)$ for $k$ gene trees. We can compute $w(U^{(g,u)})$ in $O(deg(u)^2)$. Thus, $\sum_{g \in \mathcal{G}} \sum_{u \in g} w(U^{(g,u)})$ can be computed in $O(c^2nk)$ time.

The complexity of gain computation is dominated by the gain computation of dummy taxa. We can calculate gains of all dummy taxa in $O(c^2nkd)$. An iteration of the FM algorithm consists of transferring $r$ real taxa and $d$ dummy taxa. After each transfer, the gain is recalculated. Therefore, the complexity of an iteration is $O((r+d)c^2nkd)$. The FM algorithm may take multiple iterations before finding the final bipartition where the maximum cumulative gain is non-negative. We assume that the number of iterations to find the final bipartition is bounded by a parameter $\alpha$. Empirically we have found $\alpha$ to be a very small number ($\alpha \leq 5$) even though we experimented exhaustively with a varying number of taxa, ranging from 37 to 2000, and other model parameters.

Therefore, given a bipartition, the total time complexity for improving the bipartition through the FM algorithm is $O(\alpha(r+d)c^2nkd)$.

Now, for the initial bipartition, the creation of the greedy consensus tree from the gene trees at the beginning of the algorithm requires $O(n^2k)$ time. We need to consider at most $O(n)$ edges of the consensus tree for a subproblem where we construct candidate initial bipartitions from each of the edges and calculate the score for each of them. Each score calculation step takes $O(c^2nkd)$ time. Therefore, creating an initial bipartition takes $O(c^2n^2kd)$ time in total.

Thus, handling each subproblem takes $O(c^2n^2kd + \alpha(r+d)c^2nkd)$ time. Since, at worst case $O(r+d) = O(n)$, we have the final time complexity of $O(\alpha c^2n^2kd)$ for a subproblem.

#### 3.1 Time complexity for fully resolved gene trees with the assumption of balanced partitioning

For unrooted fully resolved or binary gene trees, $c = 3$. We consider the case where each bipartition is made as evenly as possible. That is, if $|A| + |B|$ is even, then $|A| = |B|$. Otherwise, $|A| - |B| = \pm 1$. Then, the height of the subproblem tree (Example in Figure 1) is $O(\log n)$. Since a dummy taxa is introduced at each level, a subproblem can have at most $O(\log n)$ dummy taxa, that is $d = O(\log n)$. Therefore, the complexity of handling a subproblem is $O(\alpha n^2 k \log n)$. We note that each divide step in our algorithm constructs an edge of the output species tree in the corresponding conquer step. Therefore, we have $\Theta(n)$ divide steps in our algorithm. Each divide step increases the number of subproblems by 1. Therefore, we have $\Theta(n)$ subproblems and the total time complexity is $O(\alpha n^3 k \log n)$ or $O(n^3 k \log n)$ since empirically $\alpha$ is very small as mentioned before.

##### 3.2 Time complexity without the assumptions

We have proved the time complexity of $O(n^3 k \log n)$ under the assumptions of balanced bipartitions and fully resolved gene trees. Without the assumption of balanced partitioning, we obtain a complexity of $O(n^4 k)$. Finally, for gene trees with polytomies, the algorithm may experience a notable increase in running time, particularly when confronted with a large amount of polytomy. However, we do not observe a drastic rise in time in most of the cases as shown in Table 1. Nevertheless, we can readily address the issue of polytomy by resolving them and then applying wQFM-TREE on fully resolved gene trees. This approach yields equivalent performance to wQFM-TREE without resolving polytomy (Figure 4).

#### 4 Additional results

##### 4.1 Comparing wQFM-TREE with wQFM

wQFM-TREE performs the same calculations as wQFM in terms of scoring candidate bipartitions and associated steps, but directly from the gene trees by circumventing the need for explicitly generating $\Theta(n^4)$ quartets. We made an effort to make wQFM scalable without sacrificing accuracy. Nevertheless, wQFM-TREE might not be the same as wQFM in terms of the initial bipartition and the normalization process. As a result, the accuracy of wQFM-TREE may differ from wQFM. However, our findings on the 37-taxon and 48-taxon datasets presented in Fig. 3 show that wQFM-TREE is generally as good as wQFM. Out of 240 data replicates across various model conditions in the 37-taxon dataset, wQFM and wQMF-TREE generated identical trees in 238 replicates, differing in only two instances. Conversely, on the 48-taxon dataset, wQFM-TREE exhibited inferior performance in two specific model conditions, which are 0.5X-1000gt-500bp and 1X-1000gt-500bp. However, no statistically significant differences were observed between them in the remaining five model conditions.

In terms of running time, quartet generation requires a significant amount of time in the case of wQFM. On the 37-taxon dataset (1X-800gt-500bp model condition), wQFM requires approximately 140 seconds to generate quartets, followed by just 4 seconds to estimate a tree from the weighted quartets. In contrast, wQFM-TREE estimates species trees directly from the gene trees in only 4 seconds. For the 48-taxon dataset, wQFM takes about 560 seconds to generate weighted quartet distributions and an additional 14 seconds to estimate species trees from these weighted quartets. Conversely, wQFM-TREE takes 27 seconds to estimate species trees directly from the gene trees.

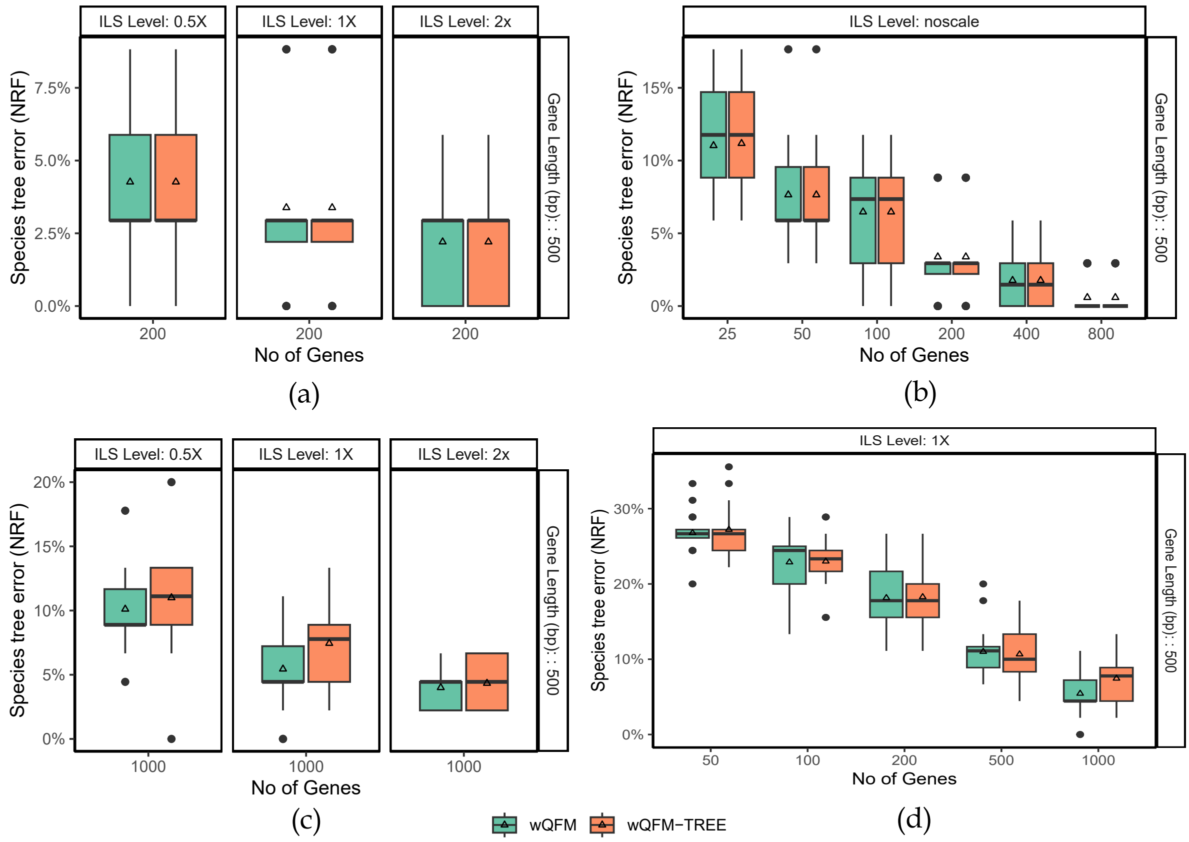

Figure 3: **Comparison of wQFM and wQFM-TREE on 37-taxon mammalian and 48-taxon avian simulated datasets.** We show the average RF rates with standard error bars over 20 replicates. (a and b) Results on the 37-taxon dataset. (a) The level of ILS was varied from 0.5X (highest) to 2X (lowest) amount, keeping the sequence length fixed at 500 bp and the number of genes at 200. (b) The number of genes was varied from 25 to 800, with 500 bp sequence length and noscale ILS. (c and d) Results on the 48-taxon dataset with varying levels of ILS and varying numbers of genes, respectively.

#### 4.2 Handling polytomies

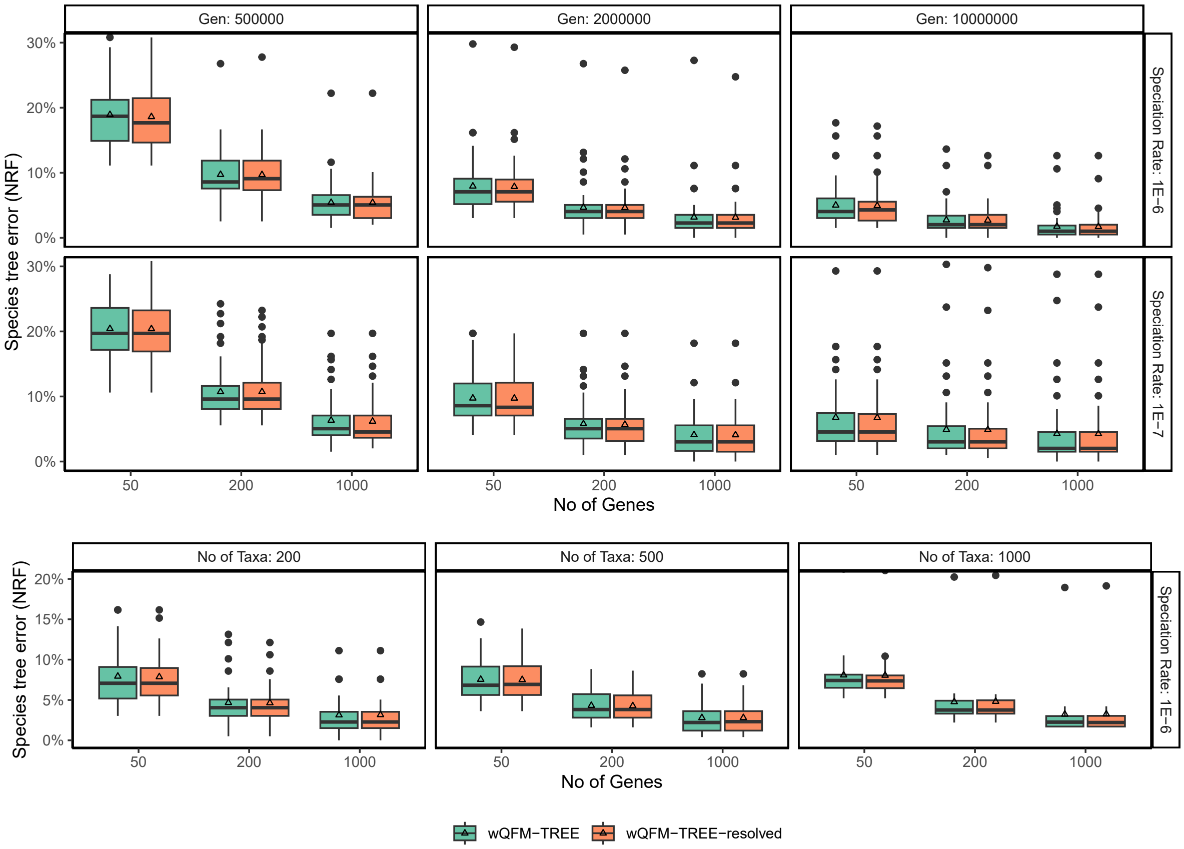

Figure 4: **Performance of wQFM-TREE with and without resolving polytomies.** (Top) Two hundred taxa and varying tree shapes and number of genes. (Bottom) Varying numbers of taxa and genes and the tree shape fixed to 2 M/1e-6.

#### 4.3 Running time comparison

Table 1: **Runtime (in seconds) of various methods on the datasets analyzed in this study.** We show the running time of wQFM-TREE with and without resolving polytomies in the gene trees. Values are shown as average (over 10 replicates for 200- and 500-taxon and 5 replicates for 1000-taxon) ± standard deviation.

| No. of taxa | No. of genes | TREE-QMC | ASTRAL | wQFM-TREE resolved | wQFM-TREE unresolved |
|---|---|---|---|---|---|
| 200 | 50 | 1.8±0.1 | 4.3±1.2 | 15.1±1.5 | 20.1±3.8 |
| 200 | 200 | 7.4±0.5 | 19.8±3.5 | 76.6±3.5 | 93.9±6.9 |
| 200 | 1000 | 37.1±1.2 | 297.6±15.8 | 391.1±11.1 | 439.5±13.4 |
| 500 | 50 | 11.4±0.6 | 24.4±3.6 | 155.3±4.6 | 156.6±5.9 |
| 500 | 200 | 43.8±1.0 | 109.1±8.9 | 584.5±10.1 | 793.5±23.4 |
| 500 | 1000 | 225.7±2.6 | 1509.4±34.4 | 2451.9±19.2 | 3413.7±45.1 |
| 1000 | 50 | 49.8±1.1 | 157.5±10.4 | 898.4±19.5 | 943.2±19. |
| 1000 | 200 | 188.0±2.3 | 658.9±22.5 | 2786.8±34.37 | 3155.8±36.9 |
| 1000 | 1000 | 913.9±5.0 | 7336.9±71.4 | 11680.4±69.9 | 12275.2±69.7 |
| 2000 | 1000 | 3394 | 2161.4 | 24552 | 24552 |
| Green plant data | | | | | |
| 1178 | 410 | 415.3 | 3340.8 | 11318.7 | 19744.8 |
